# A Cytological F1 RNAi Screen for Defects in *Drosophila melanogaster* Female Meiosis

**DOI:** 10.1101/2024.01.12.575435

**Authors:** William D. Gilliland, Dennis P. May, Amelia O. Bowen, Kelly O. Conger, Doreen Elrad, Marcin Marciniak, Sarah A. Mashburn, Gabrielle Presbitero, Lucas F. Welk

## Abstract

Genetic screens for recessive alleles induce mutations, make the mutated chromosomes homozygous, and then assay those homozygotes for the phenotype of interest. When screening for genes required for female meiosis, the phenotype of interest has typically been nondisjunction from chromosome segregation errors. As this requires that mutant females be viable and fertile, any mutants that are lethal or sterile when homozygous cannot be recovered by this approach. To overcome these limitations, our lab has screened the VALIUM22 collection produced by the Harvard TRiP Project, which contains RNAi constructs targeting genes known to be expressed in the germline in a vector optimized for germline expression. By driving RNAi with GAL4 under control of a germline-specific promoter (*nanos* or *mat-alpha4)*, we can test genes that would be lethal if knocked down in all cells, and by examining unfertilized metaphase-arrested mature oocytes, we can identify defects associated with genes whose knockdown results in sterility or causes other errors besides nondisjunction.

We screened this collection to identify genes that disrupt either of two phenotypes when knocked down: the ability of meiotic chromosomes to congress to a single mass at the end of prometaphase, and the sequestration of Mps1-GFP to ooplasmic filaments in response to hypoxia. After screening >1450 lines of the collection, we obtained multiple hits for both phenotypes, identified novel meiotic phenotypes for genes that had been previously characterized in other processes, and identified the first phenotypes to be associated with several previously uncharacterized genes.

## Introduction and Background

Genetic screens are one of the main tools for identifying which genes are required for biological processes of interest. Screens for defects in female meiosis have been conducted using mutants that were isolated from natural populations (Sandler *et al*. 1968), or induced by mutagens such as EMS (Baker and Carpenter 1972) or P element transposition (Sekelsky *et al*. 1999). These screens all made mutant chromosomes homozygous, in order to identify recessive alleles that cause females to have increased numbers of aneuploid progeny arising from meiotic nondisjunction. While effective, this criterion limits the type of mutations that might be recovered from the screen. Requiring homozygous mutant females to be viable and fertile necessarily precludes the recovery of mutants that would be lethal or cause female sterility when homozygous, and only mutants that result in nondisjunction, as opposed to other defects, will be identified. An improvement to this approach was to induce the formation of homozygous germline clones by mitotic recombination using the FLP-FRT system (Page *et al*. 2007). This has the advantage of only inducing homozygosity in the germline, while retaining heterozygosity in the soma, which permits examining the effects of mutations that are otherwise lethal as homozygotes. However, for identification, this approach still requires that the homozygous mutant clonal lineages are competent to produce viable oocytes and drive high rates of meiotic nondisjunction.

The development of tools to use RNA interference (RNAi) to knock down genes of interest in a tissue-specific manner has made it possible to circumvent some of these restrictions in genetic screens. RNAi coopts an endogenous cellular defense mechanism by introducing a palindromic sequence that is complementary to a targeted gene of interest. When transcribed, this sequence forms a double-stranded RNA hairpin, which is then processed by the cell to produce RNA-Induced Silencing Complexes that degrade the mRNA molecules of the target gene. This causes a significant knockdown in the expression levels of that gene, usually by 80-95% (Heigwer *et al*. 2018). Collections of *in vivo* RNAi constructs have been made in *Drosophila* with UAS promoters (Cook *et al*. 2010), which can then be driven by crossing to constructs expressing the yeast protein GAL4 under the transcriptional control of various *Drosophila* promoters that limit expression to specific tissues. By expressing GAL4 only in tissues that are not required for survival (such as the germline), genotypes can be assayed that would be lethal if RNAi were driven in all tissues. One complicating factor is that the germline has defense mechanisms to limit the propagation of transposable elements. This means that the original UASt vectors produced in *Drosophila* do not express well in the germline, and a different promoter, UASp, was developed that incorporates the P-element basal promoter to drive expression (Rørth 1998). This promoter was used by the Harvard TRiP project in the production of the VALIUM22 collection of RNAi stocks, which is optimized for germline expression at the cost of poorer somatic expression (Ni *et al*. 2011).

Our lab has been studying two processes that take place during Prometaphase I of female meiosis. The first is chromosome congression. *Drosophila* oogenesis has been divided into 14 stages based on morphological criteria (King 1970), and these stages track progression through the meiotic cell cycle. The stage 12/13 transition, marked by the breakdown of the nuclear envelope and the start of dorsal appendage growth, is the end of Prophase I and the beginning of Prometaphase I (Theurkauf and Hawley 1992). Once the spindle forms, the obligately nonexchange chromosome *4* (along with other nonexchange chromosomes, such as the 6-10% of spontaneously achiasmate *X* chromosomes in normal flies or 100% of achiasmate *X* chromosomes in *FM7/X* balancer heterozygotes (Zhang and Hawley 1990)) move out on to opposite sides of the meiotic spindle (Theurkauf and Hawley 1992). Despite not having chiasmata, these homologs nevertheless remain physically connected by heterochromatic tethers (Hughes *et al*. 2009) which is thought to be how the distributive system mediates the proper segregation of nonexchange chromosomes (Gilliland *et al*. 2014). The nonexchange chromosomes then undergo congression, slowly moving in and rejoining the exchange chromosomes at the metaphase plate over several hours and forming a well-defined compact karyosome at metaphase arrest (Figure 1) (Gilliland *et al*. 2009a). Meiotic nondisjunction appears to be caused by this karyosome forming with pairs of homologous chromosomes attached to the same spindle pole. This means that even though congression to a single mass completed successfully, anaphase would result in gametes receiving either 0 or 2 copies of that homolog. This malorientation explains why the numbers of nullo and diplo progeny are often equal in many mutant genotypes (Gillies *et al*. 2013).

**Figure 1:**
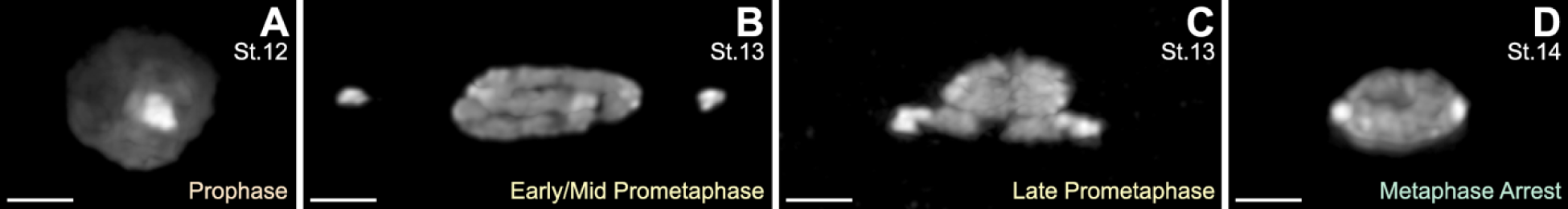
Chromosome Congression in Wildtype Oocytes. **1A)** Prophase I ends in *Drosophila melanogaster* female meiosis at the transition between oocyte stages 12 and 13, with the chromosomes in a single mass and all the heterochromatin (bright staining) clustered together. **1B)** After the germinal vesicle breaks down and the spindle forms during oocyte stage 13, nonexchange chromosomes (such as the obligately achiasmate *4* chromosomes) move out onto opposite arms of the spindle, with the exchange chromosomes remaining at the metaphase plate. **1C)** The nonexchange chromosomes then undergo congression and move in to join the other chromosomes, a process that takes several hours (Gilliland *et al*. 2009b). **1D)** Stage 14 oocytes have completed congression when the cyst arrests at metaphase, with all chromosomes in a single, well-structured mass. Note the brighter-staining heterochromatin that is clustered at each end of the karyosome. (DNA/DAPI, Scale bars = 2 µm)

We also seek to understand the composition of ooplasmic filaments, which were first identified by the localization of Mps1 and Polo proteins to them (Figure 2) (Gilliland *et al*. 2007). These filaments appear to assemble shortly after germinal vesicle breakdown (GVBD) (Gilliland *et al*. 2009b) and appear to be disassembled shortly after egg activation and fertilization (Pandey *et al*. 2007). It has been shown that the Mps1-GFP and Polo-GFP proteins reversibly localize to these filaments in response to hypoxia on a time scale of ∼10 minutes, and that a second round of hypoxia following a return to normoxia reveals the same structures, indicating that there must be a static scaffold that the Mps1-GFP or Polo-GFP proteins are being dynamically sequestered to (Gilliland *et al*. 2009b). Immunogold electron microscopy identified the scaffold as a triple-helical proteinaceous structure of ∼150 nm diameter (Gilliland *et al*. 2009b). However, despite attempted immunolocalization of many candidate cytoskeletal proteins, the identities of the proteins that build these filament scaffolds remain unknown, which has hampered efforts to determine the functional reasons for this sequestration.

**Figure 2:**
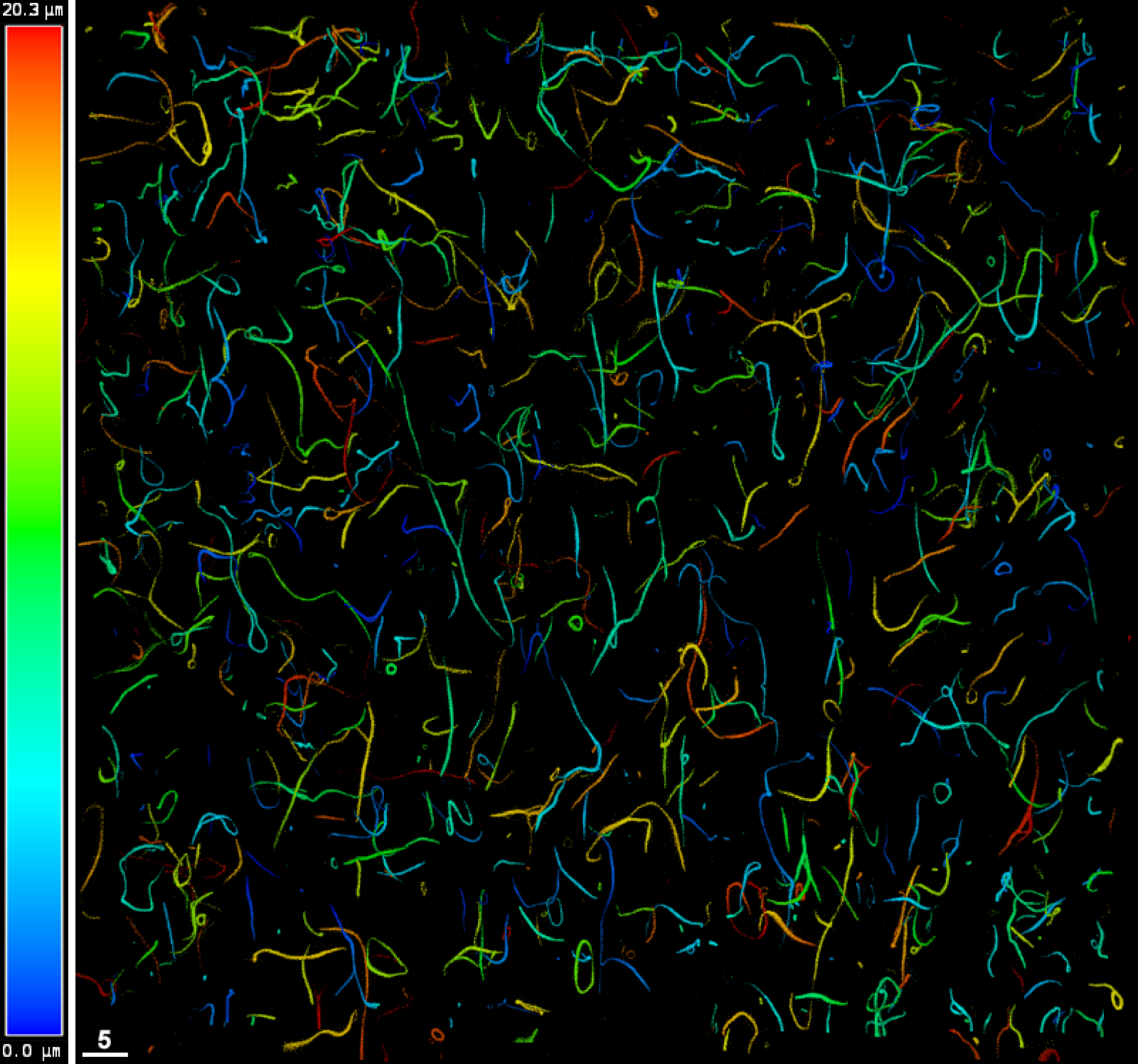
Mps1-binding Filaments. A maximum intensity projection of a 20.3 µm deep Z-stack of Mps1-GFP fluorescence from a Nanos Screen Stock oocyte that was fixed during hypoxic sequestration to the filaments. In normoxic cells, the Mps1-GFP protein is distributed diffusely throughout the oocyte. However, after exposure to hypoxia, the protein becomes reversibly sequestered to filaments distributed throughout the oocyte, on a time scale of around 10 minutes (Gilliland *et al*. 2009b). The false-color coding is for the Z-axis positioning of the filaments, which shows that most filaments begin and end within the volume (as neither end is dark red or dark blue, nor does it touch an edge), and that the GFP-labeled filaments do not form a single continuous network. (GFP, scale bar = 5 µm)

We have performed an F1 cytological screen of the VALIUM22 collection by scoring unfertilized eggs instead of adult progeny, with the goal of identifying genes that are required for these two processes but that may have been unrecoverable in previous screens due to their loss causing lethality or sterility. To conduct this screen, VALIUM22 stocks were ordered from the Bloomington Drosophila Stock Center, and males were crossed to females from a tester stock that introduced a germline-specific GAL4 driver to induce RNAi of the target, along with Mps1-GFP to allow visualization of the ooplasmic filaments. We used an optimized assembly-line protocol to dissect, fix and DAPI stain ovaries from F1 females from each cross, and mature oocytes were then visually inspected for defective phenotypes by confocal microscopy. By only driving RNAi in the germline, we could observe phenotypes for genes that would have caused lethality if eliminated from the soma, and by examining unfertilized oocytes we could not only identify defects in sterile genotypes (which would not have been possible in screening approaches that examine the progeny of experimental females) we could also identify defects that might not result in nondisjunctional progeny.

The congression defect phenotype is that the meiotic chromosomes fail to form a single mass, resulting in multiple masses at metaphase arrest. There could also be the degradation and loss of the meiotic chromosomes, or other obvious defects in the structure of the compact karyosome, such as all the heterochromatin being in a single mass instead of two masses at opposite ends of the karyosome. The phenotype of defects in the filaments is a complete lack of sequestration during hypoxia, resulting in no detectable filaments, or other gross changes to the morphology of the filaments. Because a previous RNAi screen looking for defects in germline stem cell maintenance found a number of genes that resulted in a complete oogenesis failure after knockdown (Yan *et al*. 2014), we expected that some lines would fail to produce mature stage 14 oocytes to test when driven by *nos::Gal4*. We therefore retested any lines that did not produce mature oocytes in the initial cross by using a second driver stock carrying *matα4::GAL4,* which turns on later in oogenesis.

We screened 1,461 constructs that targeted different genes. While a number of these genes were already known to be required for meiosis, we did not want to exclude any genes from evaluation because most of these genes have never been specifically examined for either phenotype, and known meiotic mutants could serve as internal controls (e.g. in the pilot project for the screen, a construct targeting Mps1 was added anonymously to the lines being tested. This line was successfully identified as having no visible filaments as well as no congression defect, results consistent with both successful RNAi knockdown of the Mps1-GFP reporter gene as well as the known congression phenotype seen in mutant *mps1* genotypes (Fischer *et al*. 2004; Gillies *et al*. 2013)). Of those constructs tested with *nos::Gal4*, we found 1,265 constructs with negative results, 156 that produced no mature oocytes, and 38 that were hits. When the 156 constructs were retested with *matα4::GAL4,* we found that 32 produced no mature oocytes and so were untestable in this assay, 108 constructs had no phenotype and were designated “*nos* duds”, and 16 were hits with phenotypes of interest, for a total hit rate of 3.7% (54/1461) for the entire screen. Over two-thirds of our hits were sterile or semi-sterile when undergoing RNAi, confirming that our screening approach could successfully identify genes of interest that would likely not have been recoverable using traditional meiotic screening methods.

## Methods

### Screen Stocks

The primary Nanos Screen Stock (NSS) was *FM7w, y w v B; P{mps1-GFP, w^+^}II.1, P{nos::Gal4, w^+^}; sv^spa-pol^,* while the secondary Mat Alpha Screen Stock (MASS) was *FM7w, y w v B / y w; P{mps1-GFP, w^+^}II.1 P{mat-α4-GAL-VP16, w^+^}V2H / CyO; sv^spa-pol^,* with males of both stocks also carrying a labeled *y^+^Y* chromosome. The second chromosome in MASS was created by crossing the *P{mat-α4-GAL-VP16}V2H* construct from Bloomington stock 7062 to the *P{mps1-GFP}II.1* chromosome and selecting for recombinant progeny with a more saturated orange eye color than either single construct. This recombinant chromosome was then crossed into the same *FM7w, y w v B; sv^spa-pol^* background as NSS, followed by validation that both P elements being present by examining GFP fluorescence and the ability to drive RNAi. While the NSS was homozygous viable, after multiple attempts to isogenize the MASS we found that females were only weakly fertile when homozygous for either the *FM7* or *P{mps1-GFP, w^+^}II.1 P{mat-α4-GAL-VP16, w^+^}V2H* chromosomes. This was unexpected, as both chromosomes are fertile when homozygous on their own. Therefore, the MASS had to be maintained by manually selecting for *Bar* and *Curly* males and females when doing stock transfers.

### Screen Approach

Lines were tested through a standard pipeline (Figure 3). VALIUM22 lines were ordered from BDSC every two weeks in blocks of 40-50 lines, in a randomized order. Stocks arrived labeled only by their BDSC stock number, without reference to the identity of the gene targeted by the hairpin construct. The week the shipment arrived, males were collected from half of the vials and crossed to virgin NSS females in yeasted vials; males from the other half of the vials were similarly crossed the following week. After laying eggs for one week, adults were removed from the cross vials, and starting on day 10, up to 10 F1 females (*FM7w, y w v B / y v sc; P{mps1-GFP, w^+^}II.1 P{mat-α4-GAL-VP16, w^+^}V2H / P{UAS-RNAi, y^+^ v^+^}; sv^spa-pol^/+* for RNAi constructs inserted on chromosome II, or *FM7 / y v sc; P{mps1-GFP, w^+^}II.1 P{mat-α4-GAL-VP16, w^+^}V2H / +; P{UAS-RNAi, y^+^ v^+^}/+; sv^spa-pol^/+* for RNAi constructs inserted on chromosome III) were collected from the progeny of each vial, then aged 3-4 days in yeasted vials to prepare for dissection. After dissection and fixation, the slides were scored the following week. If a line was found to not be a hit, the stock vial was discarded. This approach saved resources, as most stocks could be tested before the shipped vial stock needed to be transferred to fresh media. When a line crossed to NSS produced F1 females with no mature oocytes, that line was subsequently crossed to MASS virgin females, to collect 10 F1 females *(FM7w, y w v B / y v sc; P{mps1-GFP, w^+^}II.1 P{mat-α4-GAL-VP16, w^+^}V2H / P{UAS-RNAi, y^+^ v^+^}; sv^spa-pol^/+*) for dissection. If MASS-crossed F1 females still produced no mature oocytes, the line was categorized as untestable in our assay and discarded. If instead the MASS-crossed F1 females produced mature oocytes, it was classified as a hit if those oocytes had phenotypes of interest, otherwise it was classified as a *nos* dud and discarded. For nine RNAi stocks, no F1 progeny were produced when crossed to tester stock females. Reasoning that this could be due to maternally loaded GAL4 protein causing early embryonic lethality, these stocks were crossed in the other direction (RNAi-stock virgin females mated to NSS or MASS males), which did produce living F1 females in all cases, and resulted in 4 negative results and 5 *nos* duds.

**Figure 3:**
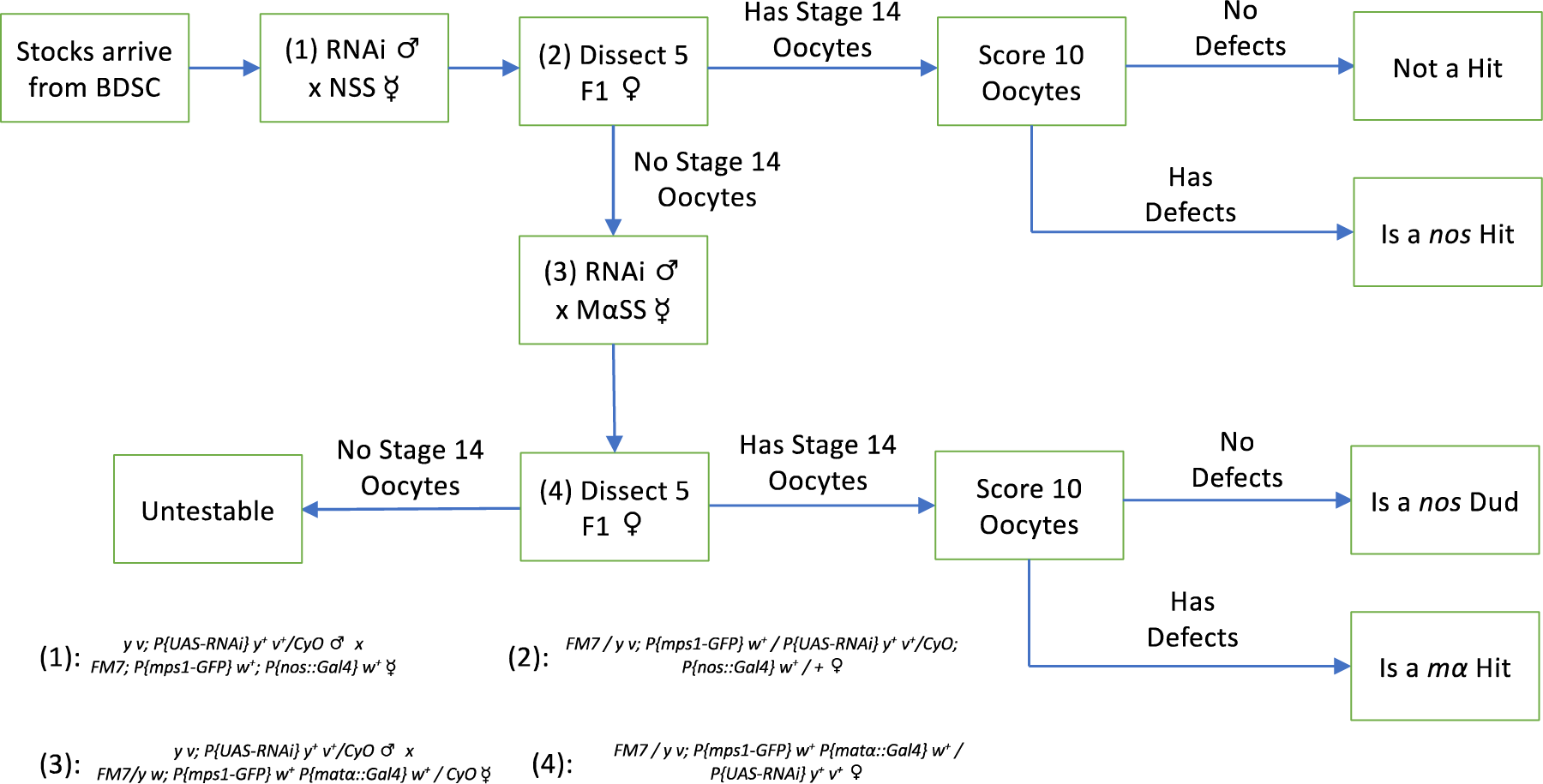
Screen Crossing Pipeline. A schematic showing how lines were crossed to test for meiotic defects. Males from newly-arrived RNAi stocks were crossed to NSS virgin females (cross genotype 1), and F1 females (genotype 2) were dissected. If these females had mature oocytes, they were scored on the microscope and the line classified as a hit or not. However, if (2) females had no oocytes, males from the RNAi stock were crossed to MASS virgin females (cross genotype 3) and F1 females (genotype 4) were dissected. If (4) females had mature oocytes, the line was scored and classified as either a hit or a *nos* dud, but if oocytes were present the line was classified as untestable. This diagram is for RNAi constructs on chromosome *2*; the same steps (with slightly different genotypes) were done for constructs on *3*.

### Streamlined Screen Dissection with Hypoxia

To facilitate the large numbers of dissections required for this project, we streamlined our standard ovary prep (Gillies *et al*. 2013) to reduce the time required, mainly by reducing the number of wash steps. Starting at time zero, we dissected five heterozygous females in 1x Robb’s media (Sullivan *et al*. 2000) + 0.1% BSA + 0.03% sodium azide. (The azide inhibits mitochondrial respiration, which enhances the hypoxic response and yields better localization of Mps1-GFP to the filaments (Gilliland *et al*. 2009b)). The tube was closed until 30 minutes had elapsed, then the Robb’s was aspirated off and 1 ml of fixative, a freshly combined 1:1 mix of 16% EM grade paraformaldehyde (Ted Pella) and 2x WHOoPASS buffer (Gillies *et al*. 2013) was applied, and the tube was incubated on a nutator for 6 minutes. The fixative was then replaced with PBST (PBS + 1% Triton-X 100), and the ovarioles were separated by rapidly pipetting up and down through a P1000 pipette. Oocytes were then washed for 15 minutes in 1 ml of PBST two times, then 2 µl of 500x DAPI was added to the PBST and incubated for 6 minutes, then washed in fresh PBST for 15 minutes before all oocytes were mounted on a single slide with either SlowFade Gold or SlowFade Diamond mountant (ThermoFisher), with the coverslip sealed with nail polish. To utilize this streamlined protocol in an assembly line, one person would dissect a stock of flies every 4 minutes, with the tubes being passed to others to do the washing, staining and mounting steps at the appropriate times, which allowed us to do 20-25 preps per weekly session.

In March 2020, when the screen was around 90% complete, the COVID-19 quarantine restrictions were imposed, which prevented us from working as a group in the lab. To finish the screen, the protocol was modified so a single worker (WDG) could complete 15 lines per week in two blocks of 7 or 8 lines, doing dissections every 4 minutes during the initial 30 minute incubation, then switching to fixing and washing steps for the remaining lines in the block. Once preps reached the first PBST wash, they were left on the nutator in the dark at room temperature while the second block was processed up to the same stage. Then all remaining washing, staining and mounting steps were performed with all preps in a single block.

### Slide Maps and Scoring

To facilitate efficient scoring of oocytes and avoid double-counting the same cells, a Nikon SMZ1500 dissecting microscope equipped with a DS-Fi1 camera was used to take an image of each slide suspended over a black background and illuminated from the sides with fiber optic lights using Nikon Elements D software. These slide maps were color-inverted and printed on the back of each score sheet along with the line number, and were used to facilitate locating oocytes to score.

To score a slide, the slide map was used to find 10 mature oocytes (those with fully formed dorsal appendages) on a Leica SPE II confocal microscope using the 20x objective, with the coordinates of each recorded using the LAS AF (or later LAS X) software package’s Mark and Find panel (Leica). The microscope was then switched to the 63x objective, and the Mark and Find panel was used to quickly navigate to the marked oocytes. Each oocyte was then visually scored for successful congression (having all of chromosomes together in a single mass), and whether there were visible GFP filaments present in the cell. Images were only collected for oocytes with defects, and data was recorded on the score sheet printed on the back of the slide map. All presented images were deconvolved using Huygens Essential using all default parameters, except for mounting medium refractive index, which was set to 1.42 per manufacturer’s specification.

### FISH and Antibody Ovary Preps

Further characterization of some hits involved Fluorescent *in situ* Hybridization to identify specific chromosome locations, or immunofluorescent preps to localize Mps1, tubulin or actin proteins, which were done as previously described (Gillies *et al*. 2013) using guinea pig anti-Mps1 used at 1:500 (Gilliland *et al*. 2007), rat anti-tubulin used at 1:250 (Serotec MCA786), or rhodamine phalloidin used at 1:250 (a generous gift from Dr. Elizabeth Leclair) and goat anti-rodent secondary antibodies (Invitrogen) used at 1:250.

### Fertility testing

Multiple females of the same F1 genotypes used for dissection were mated in a single vial to males from a standard nondisjunction tester stock, *C(1;Y), v f B / Ø; C(4)RM, ci ey^R^.* This cross allowed the unambiguous identification of nullo-*X* males *(w^+^ f^−^)* and nullo-4 progeny *(ci ey^R^)*. However, because the F1 females were not homozygous for *spa^pol^,* it was not possible to identify any diplo-*4* progeny produced, and because the RNAi constructs were marked with *y^+^,* half of any diplo-X females (which are *y^−^)* would also not be identifiable. Analogously to correcting for the 50% lethality of *X* nondisjunctional progeny due to fertilization by sperm with the wrong *X* genotype (Cooper 1948; Zeng *et al*. 2010), we corrected the rate of *4* NDJ by doubling the number of nullo-*4* progeny, doubling the number of nullo-*X* progeny, and quadrupling the number of diplo-*X* progeny observed (Explicitly assuming that diplo and nullo progeny are equally frequent, we double the number of diplo-*X* progeny once for the 50% lethality and again for the 50% of *y^−^* diplo-*X* females that are expected to be masked by receiving the *y^+^* RNAi construct.)

### Recombination testing

Measuring recombination was complicated by the VALIUM22 and GAL4 plasmids being located on the *2* and *3* autosomes, while the visible markers in those plasmids (*y^+^ v^+^* for VALIUM22, and *w^+^* for the GAL4) are *X*-linked loci often used for measuring recombination on the *X*. To measure recombination, we created females that were undergoing RNAi while heterozygous for the X-linked markers *cv* and *f*, which defines an interval of 43 cM between the markers according to FlyBase (Marygold *et al*. 2013) and crossed to *y cv v f* males; recombinant progeny were mutant for only one marker (either *cv f^+^ or cv^+^ f),* while nonrecombinant progeny were either wildtype or mutant for both markers. Map distance was then calculated as the fraction of recombinant progeny out of the total.

### Gene Ontology Analysis

Gene Ontology analysis was performed using the ShinyGO (Ge *et al*. 2020) server http://bioinformatics.sdstate.edu/go/ version 0.7.7 on 1/4/24. The list of flybase gene identifiers (FBgn) for the entire list of tested constructs was used as the background, and lists of FBgns for the untestable genes, *nos* duds, and screen hits were analyzed separately using the GO Biological Process pathway database with default parameters. When an RNAi construct could target two overlapping genes, both were included as separate entries, whereas constructs that targeted genes with many copies (e.g. His1) only included the first gene listed. Nodes in the resulting network graphs were manually repositioned to improve legibility of the graph.

## Results

We examined over 1450 RNAi constructs from the VALIUM22 collection by crossing each line to our screen tester stocks, which introduced three constructs to each line (Figure 3). The first construct was the *X* chromosome balancer *FM7,* which completely suppresses recombination when heterozygous over a normal sequence chromosome and forces the *X* chromosomes into the distributive segregation system; this is known to make meiosis more sensitive to disruption (Zwick *et al*. 1999). The second construct was a Mps1-GFP construct under its native promoter (Fischer *et al*. 2004), to allow us to visualize the ooplasmic filaments that the Mps1 protein sequesters to in response to hypoxia. The third construct was a germline-specific GAL4 driver, to cause transcription of the shRNA construct and induce RNA interference of the targeted gene. All lines were tested with *nos::GAL4*, while those lines that did not produce testable oocytes in that cross were subsequently tested with *matα4::GAL4*. F1 females were aged 3-4 days post eclosion, and then ovaries from five females were dissected, incubated in a sealed Eppendorf tube for 30 minutes to induce hypoxia, fixed, DAPI stained and mounted. Ten oocytes from each slide were then examined on the confocal microscope for any defects in chromosome congression and in the Mps1-binding filaments.

Our criterion for identifying screen hits with congression defects was based on the normal phenotype of the meiotic karyosome at metaphase arrest, which is to form a compact lemon-shaped mass with the heterochromatin clustered at each end (Figure 1D). When congression fails, the chromosomes will form multiple separated masses at metaphase arrest, as can be seen for example in *nod* mutants (Gillies *et al*. 2013). Because congression failure does happen spontaneously at a low rate in wildtype oocytes, we considered a line to be a hit if at least 2 out of the 10 scored oocytes contained multiple DNA masses. This threshold is based on the Binomial Probability Formula,

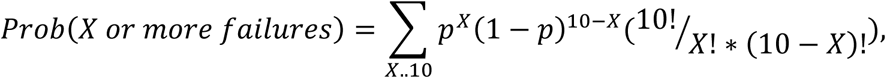

where *p* is the probability of spontaneous congression failure. If *p* is set equal to the spontaneous nondisjunction rate of 1% in wildtype *FM7/X* females (Zhang and Hawley 1990), then using a threshold of 2/10 oocytes should result in a 0.2% rate of false positives. We note that mutants in some meiotic genes, such as *mps1*, are known to cause nondisjunction without congression failure by forming a single chromosome mass that contains misaligned chromosomes (Gillies *et al*. 2013), a defect that would not be identified by this approach.

Our criteria for identifying screen hits with defects in the Mps1-binding filaments (Figure 2) was more qualitative than for congression, as the appearance of the Mps1-binding filaments depended on the hypoxic state of the oocyte at the exact moment of fixation, which could result in variation even within a single prep. To be conservative, a line was classified as normal if GFP filaments could be clearly seen in at least some of the oocytes, even if the localization to the filaments was weak, as that would indicate that genotype was still competent to construct the filaments and that Mps1-GFP was still able to localize to them. However, we also classified some lines as hits because they resulted in obvious changes to the shape of the filaments, which could be seen across multiple oocytes. We recognize that this criterion almost certainly missed genes that produced subtle changes in the filaments when knocked down, but as our primary objective was to identify the structural components that build the filaments, we reasoned that the ablation of those critical components should cause significant large-scale changes.

### The phenotypes of the screened lines could be broadly divided into four categories

#### Negative Results

We identified 1,265 lines (86.5% of the collection) that produced scoreable oocytes after RNAi knockdown was driven by *nos::GAL4*, and the mature oocytes they produced did not appear to have defects in either screened-for phenotype. The genes targeted by these constructs are listed in Table S1.

#### Untestable Genes

We identified 32 lines (2.2% of the collection) that could not be tested in our assay, as RNAi knockdown caused them to produce no scorable mature oocytes when driven by either *nos::GAL4* or *matα4::GAL4.* This included six genes that have not previously been characterized. The genes targeted by these constructs are listed in Table S2.

#### *Nos* Duds

We identified 108 lines (7.4% of the collection) that did not produce any scorable mature oocytes when driven by *nos::GAL4,* but did produce mature oocytes when driven by *matα4::GAL4*, and those oocytes were not found to exhibit defects in either screened-for phenotype. This included 16 genes that have not previously been characterized. The genes targeted by these constructs are listed in Table S3.

#### Screen Hits

We identified 38 lines (2.6% of the collection) that produced scorable oocytes when driven by *nos::GAL4,* and those oocytes exhibited defects. This included three genes that have not been previously characterized. We also identified 16 lines (1.1% of the collection) that did not produce scorable oocytes when driven by *nos::GAL4,* but did produce scorable oocytes when driven by *matα4::GAL4*, and those oocytes exhibited defects in oogenesis. This included two genes that have not been previously characterized. The genes targeted by these constructs are listed in Table 1. Overall, we found a total of 54 hits, for a hit rate of 3.7% (54/1461). While this hit rate might seem high in comparison to traditional mutagenesis screens like EMS, remember the genes included in the VALIUM22 library were selected because of their known expression in the germline, so this set of genes should be enriched for genes of interest compared to the entire genome.

**Table 1:**
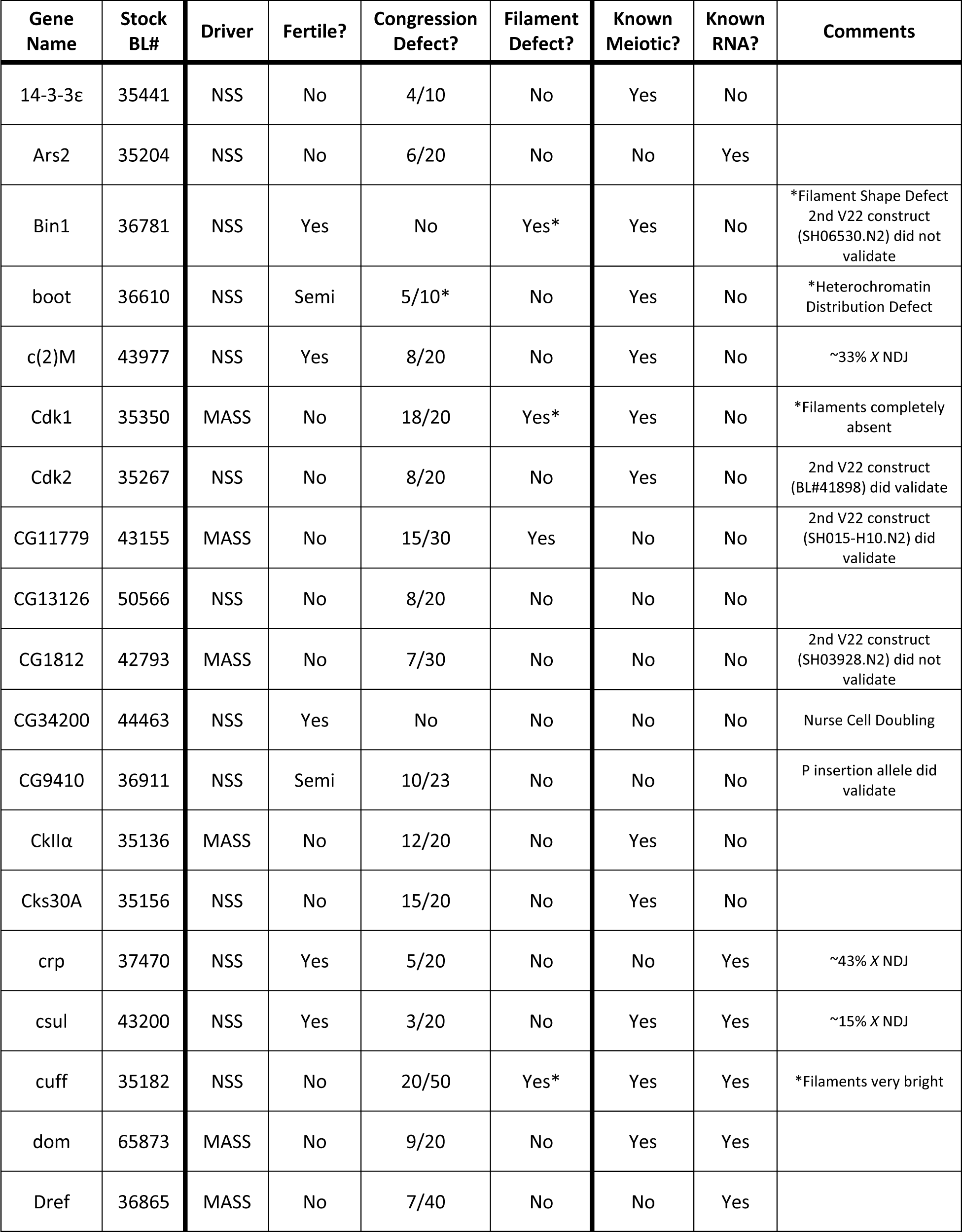
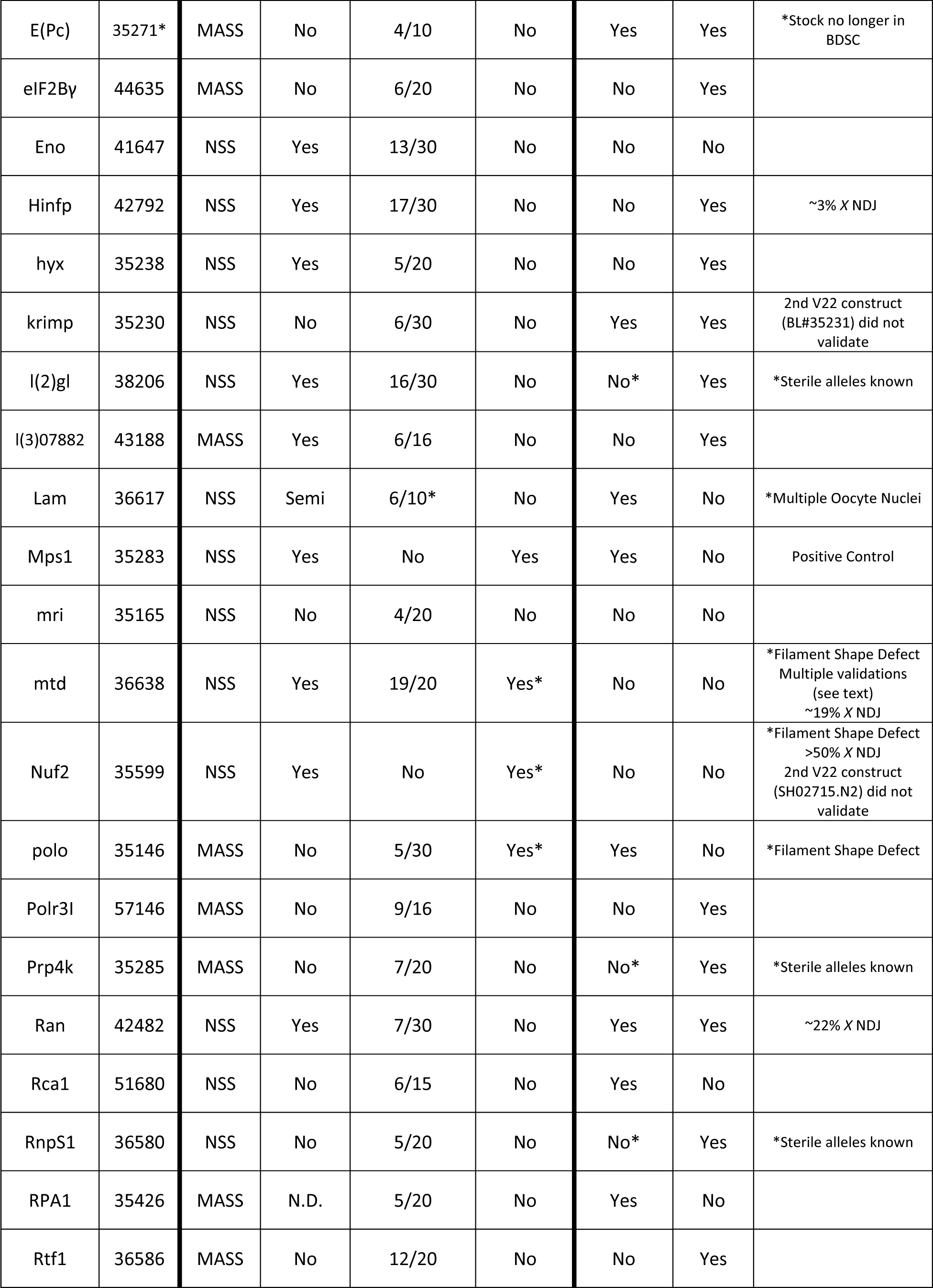
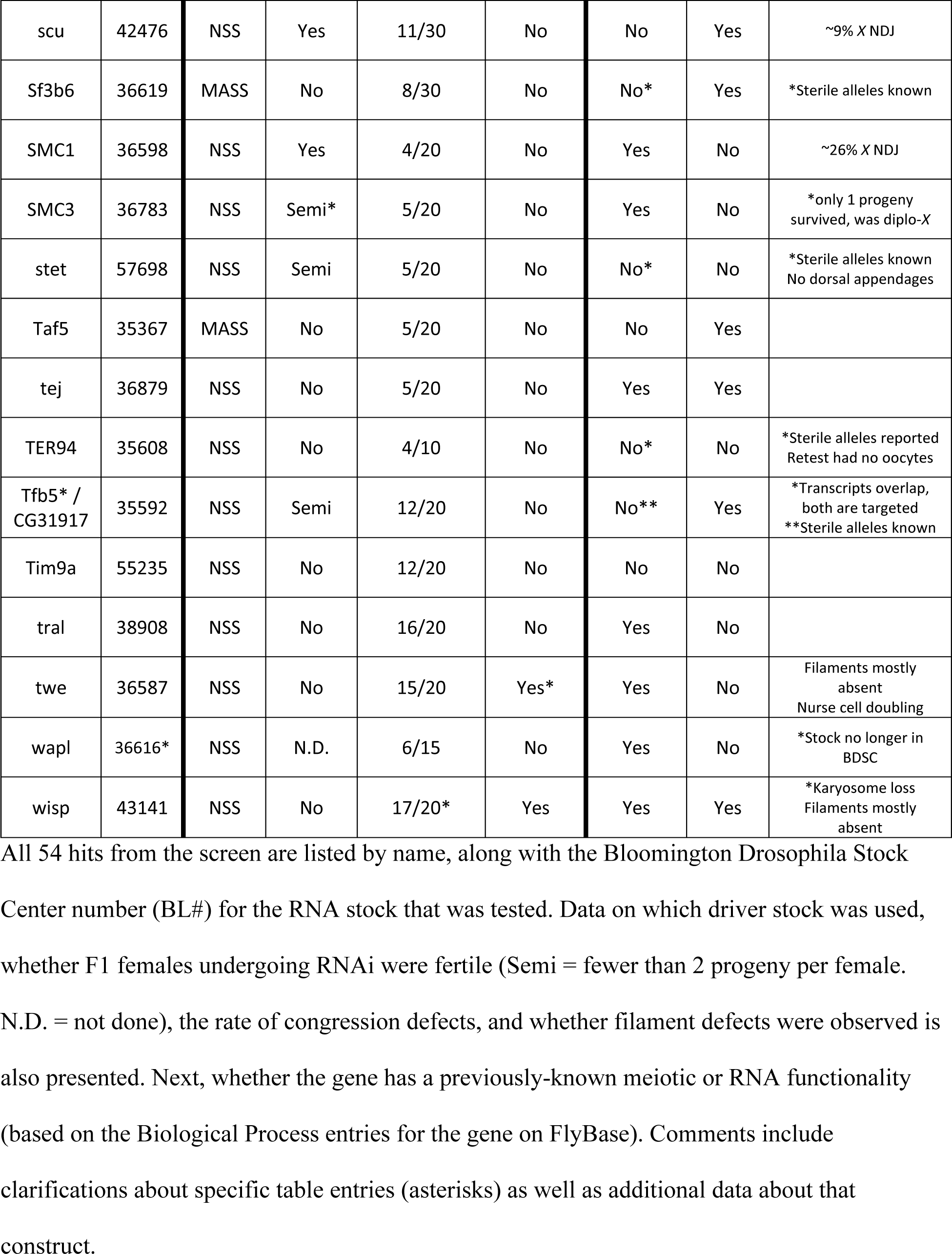
Screen Hits.

### Validation

It is important to confirm RNAi results, as nonspecific binding of the recognition sequence can lead to off-target effects (Heigwer *et al*. 2018). Our original plan was to validate screen hits using second-site RNAi constructs with recognition sequences that match different regions of the same gene. Observation of a similar phenotype with non-overlapping RNAi constructs provides strong confirmation for the effect being caused by knockdown of the gene that the construct was intended to target, as it is extremely unlikely that two entirely different sequences would cause similar off-target phenotypes. However, it is important to keep in mind that if a second-site RNAi construct fails to recapitulate what was seen with the first construct, that is a negative inference and does not necessarily mean the result of the first construct is invalid. The second construct might target a different splicing isoform (that is not required for the phenotype), or the second construct may not cause sufficient knockdown to induce the same defect. We expected this validation approach would be possible in most cases, as examination of a small random sample of genes in the VALIUM22 collection found that ∼80% had other RNAi constructs available, with some also in the VALIUM22 collection, but more in the much larger VALIUM20 collection. The VALIUM20 collection was designed to be capable of expression in both germline and somatic tissues, and RNAi knockdown by VALIUM20 constructs in the germline was reported to be “potent” (Ni *et al*. 2011). However, we found that in our assay none of the second-site constructs from the VALIUM20 vector that we tested replicated the original phenotypes, even for constructs that targeted known meiotic mutants. We believe that this is because VALIUM20 does not cause knockdown as efficiently as VALIUM22 does in the mature stage 14 oocytes we scored. We note the original testing of VALIUM20 performed by Ni *et al*. used the strong Maternal Triple Driver GAL4 stock (Mazzalupo and Cooley 2006), and they also tested genes (*bam, otu*) that were already known to cause severe defects in early stages of oogenesis. However, using our single driver stocks and examining phenotypes in mature stage 14 oocytes, we did not see any phenotypes in any of the eight VALIUM20 constructs that we tried to validate screen hits with. Because our lab does not have the resources to pursue other more costly and labor-intensive methods of validation for all hits (e.g. using CRISPr to induce silent mutations that produce RNAi-insensitive alleles), we have been unable to validate many of our screen hits, and we note that unvalidated results should be considered preliminary until the phenotypes can be confirmed using independent reagents. However, there are reasons to expect that most of the hits we identified will turn out to be valid. First, the short hairpin RNAs used to build the VALIUM22 collection were specifically designed to minimize off-target effects, by searching the rest of the genome for potential binding sites during their design (Ni *et al*. 2011). Second, nearly half (25/54) of the screen hits were to previously-identified meiotic genes, and the RNAi phenotypes we observed for many of those were entirely consistent with their known functions. Third, a number of hits were to multiple genes within similar pathways, which is consistent with it being the disruption of that process leading to the phenotypes that we observed, rather than random off-target effects.

An additional challenge for validation was that there could be variation in how effective the RNAi was, even between the two ovaries within a single female. This worked to our advantage in some circumstances; some genotypes produced only 1 or 2 small ovaries among the 5 females that were dissected, but those ovaries yielded enough mature oocytes to allow the line to be tested and reveal defects. Presumably this was because there is a threshold level of gene function required to complete oogenesis, and those cells that reached maturity happened by chance to have insufficient knockdown to fall below that threshold. However, this variation could also complicate classification. For example, TER94 was a NSS hit that produced only a few very small ovaries with a 40% congression failure rate, but when it was retested the same line did not produce any oocytes at all, which would classify it as a *nos* dud instead of a hit. To deal with discrepancies like this, we have tried to assign genes to the most appropriate category based on all of the available data, but recognize that this cell-to-cell variation means that some lines might have been classified differently under different random outcomes.

### Gene Ontology Analysis

To determine what categories of functions were associated with the genes we identified in this project, we used ShinyGO (Ge *et al*. 2020) to perform gene ontology analyses on three sets of our results: the untestable genes, the *nos* duds, and the screen hits, using the complete list of tested constructs as the background (all identifiers in Supplemental File S2). These analyses revealed that the three sets of genes largely encompassed different processes. The untestable genes (Figure 4A) were mostly in metabolic, catabolic or translational pathways, but there was relatively little involvement of meiosis-specific pathways. The *nos* duds (Figure 4B) also included genes in RNA metabolic processes, including RNA processing, RNA metabolism and ribosome biogenesis pathways, along with cell cycle, cell division and stem cell processes. In contrast, the screen hits (Figure 4C) were mostly in genes involved in mitotic and meiotic cell cycles, organelle organization, and meiotic division processes, along with a non-overlapping network of genes required for RNA processing.

**Figure 4:**
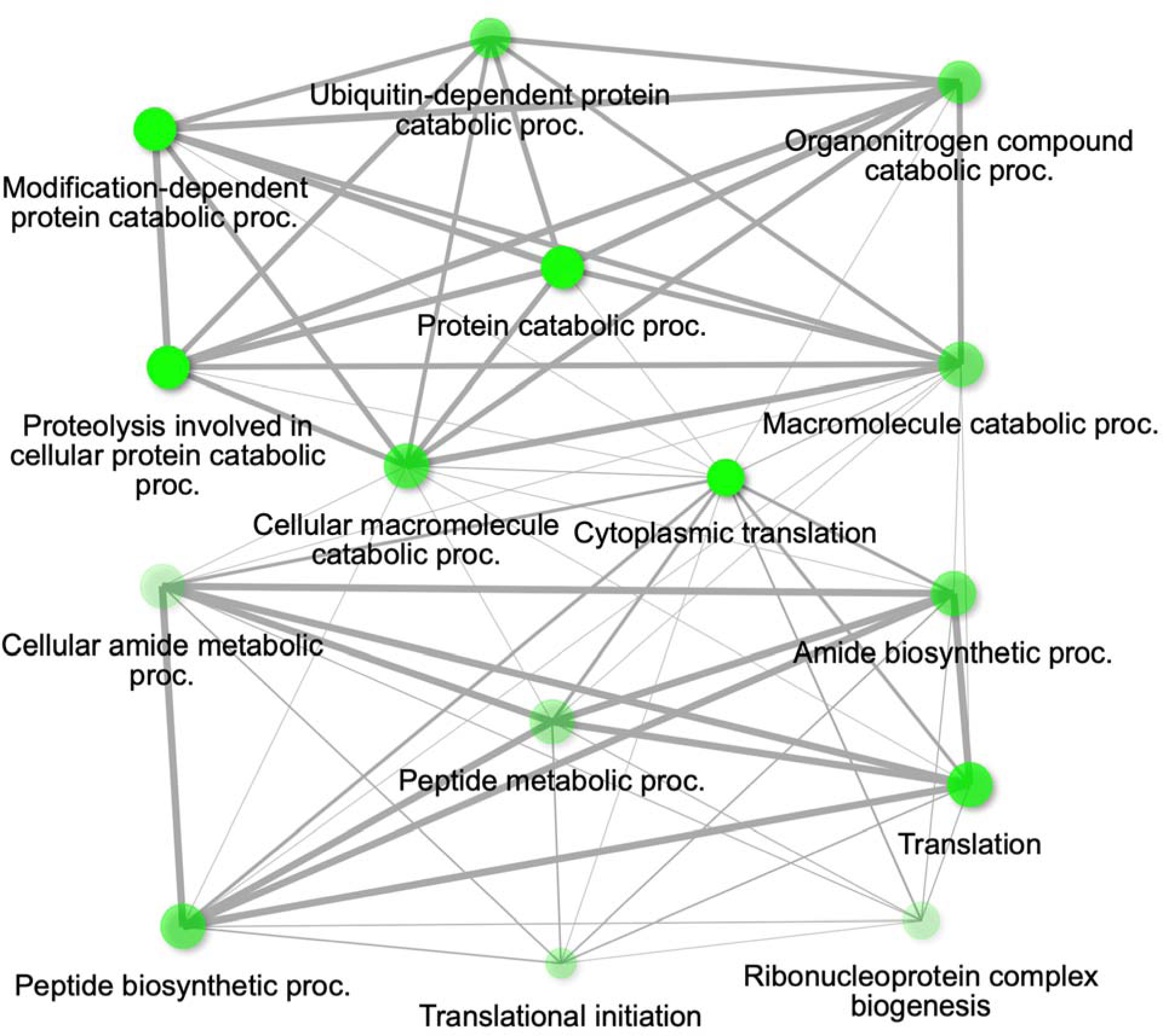
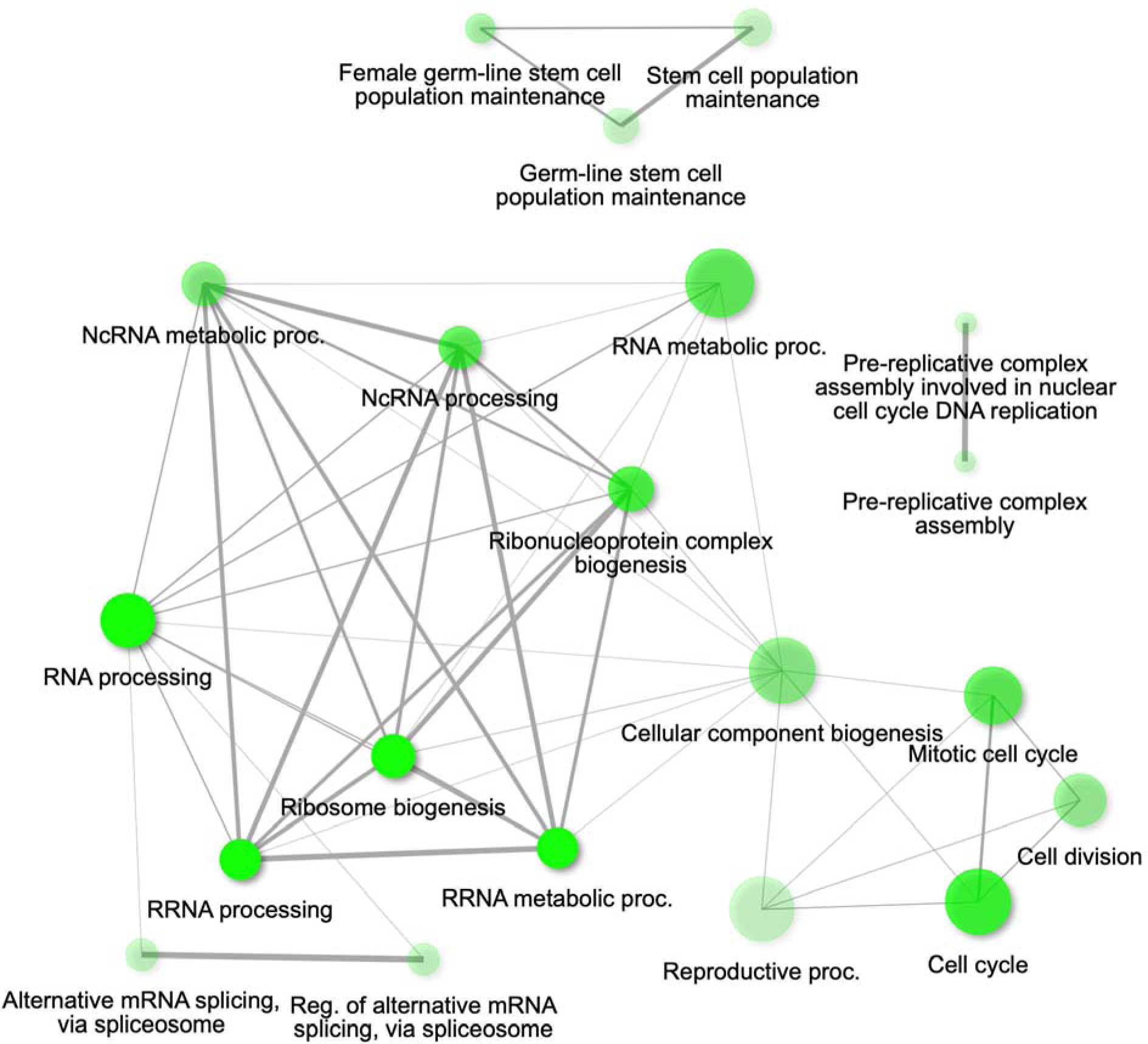
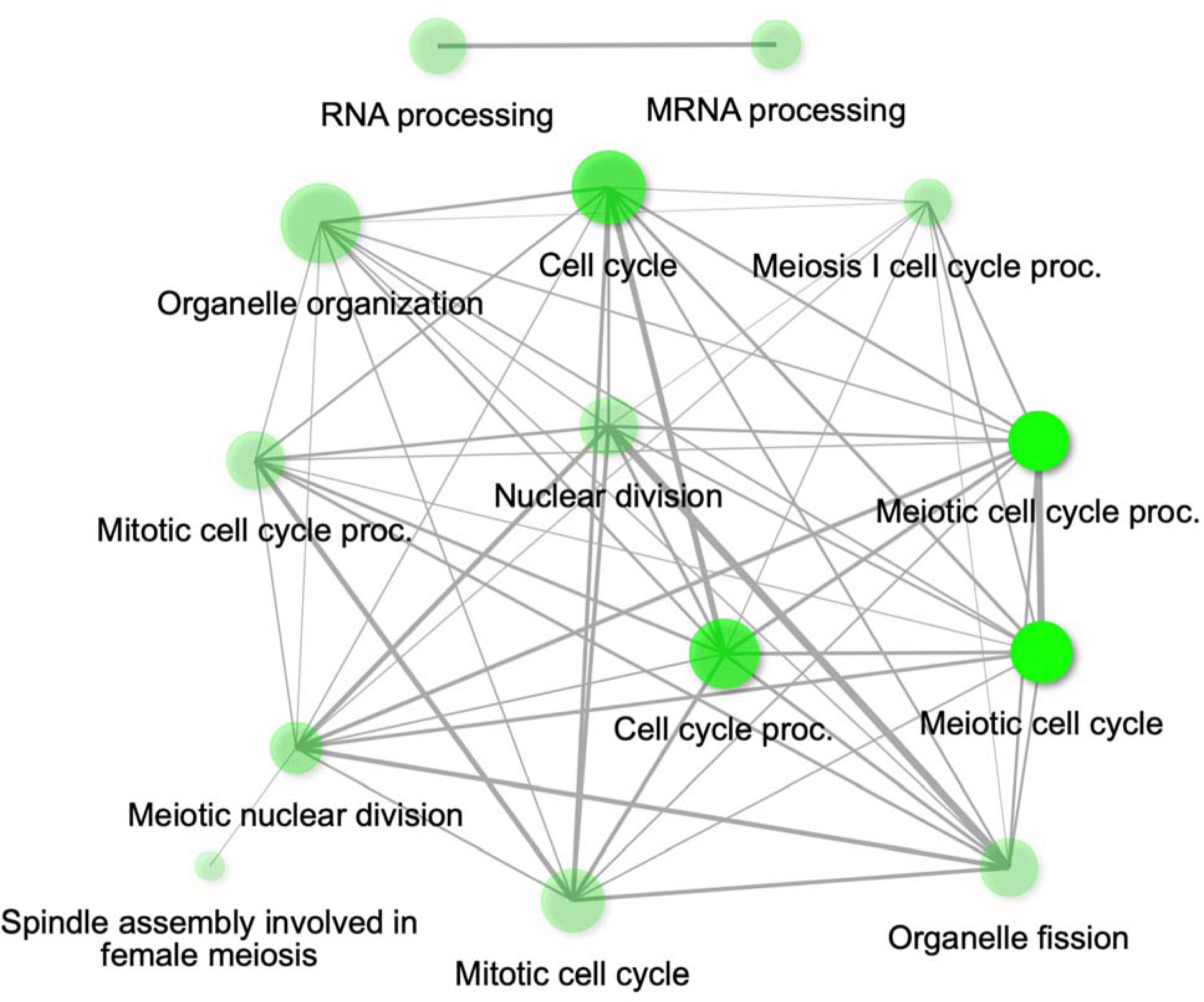
Gene Ontology Network of Untestable Genes. **4A)** The GO network diagram for the untestable genes, showing that most terms are associated with metabolic processes. The full list of genes in each node is available in Supplemental File S2. **4B)** The GO network diagram for the *nos* dud genes, showing that a number of RNA processing terms are represented, along with DNA replication, stem cell, and cell cycle processes. The full list of genes in each node is available in Supplemental File S2. **4C)** The GO network diagram for the screen hits, showing that most terms are associated with cell cycle processes, along with a separate network of RNA processing genes. The full list of genes in each node is available in Supplemental File S2.

### Description of Screen Hits

All constructs that were identified as screen hits are listed in Table 1. We have attempted validation of some of these constructs, either by testing additional RNAi constructs or by testing mutant alleles that were available from the stock center. However, because of the large number of screen hits, we have focused our efforts on those that appeared the most promising or interesting targets for further investigation.

### Screen Hits: Previously Uncharacterized Genes

Some of the most desirable results of a genetic screen are to find hits in novel genes that have never previously been characterized. In *Drosophila,* uncharacterized genes are referenced by the *CGxxx* identifiers assigned by gene prediction programs, and get assigned a name once a paper is published associating a phenotype with the gene. In addition to identifying 6 uncharacterized genes among the untestable constructs, and 16 uncharacterized genes among the *nos* duds, we had 7 uncharacterized genes among the screen hits, three of which have already been characterized in published papers before the completion of this manuscript.

#### CG1812

We identified *CG1812* as a *mat-alpha* hit with congression defects, with females undergoing RNAi producing 7/20 oocytes having multiple masses, while no defects in the Mps1-binding filaments were observed (Figure 5A). No mature oocytes were produced in females where RNAi was driven by *nos::Gal4.* A second-site RNAi construct in VALIUM22 (SH03928.N2) was created by the TRiP project. While females undergoing *matα4::Gal4* driven RNAi using the original allele were sterile, the second-site construct was found to be fertile when driven by *matα4::Gal4* and we observed no oocyte defects in that genotype. We retested the new construct using *nos::Gal4,* and again found no defects in those females. According to its entry in FlyBase (http://flybase.org/reports/FBgn0031119), *CG1812* is predicted to be a protein ubiquitinase that is orthologous to the human *kelch like protein 26 (KLHL26).* Because we have not been able to validate *CG1812* with an independent reagent, we have not named the gene at this time.

**Figure 5:**
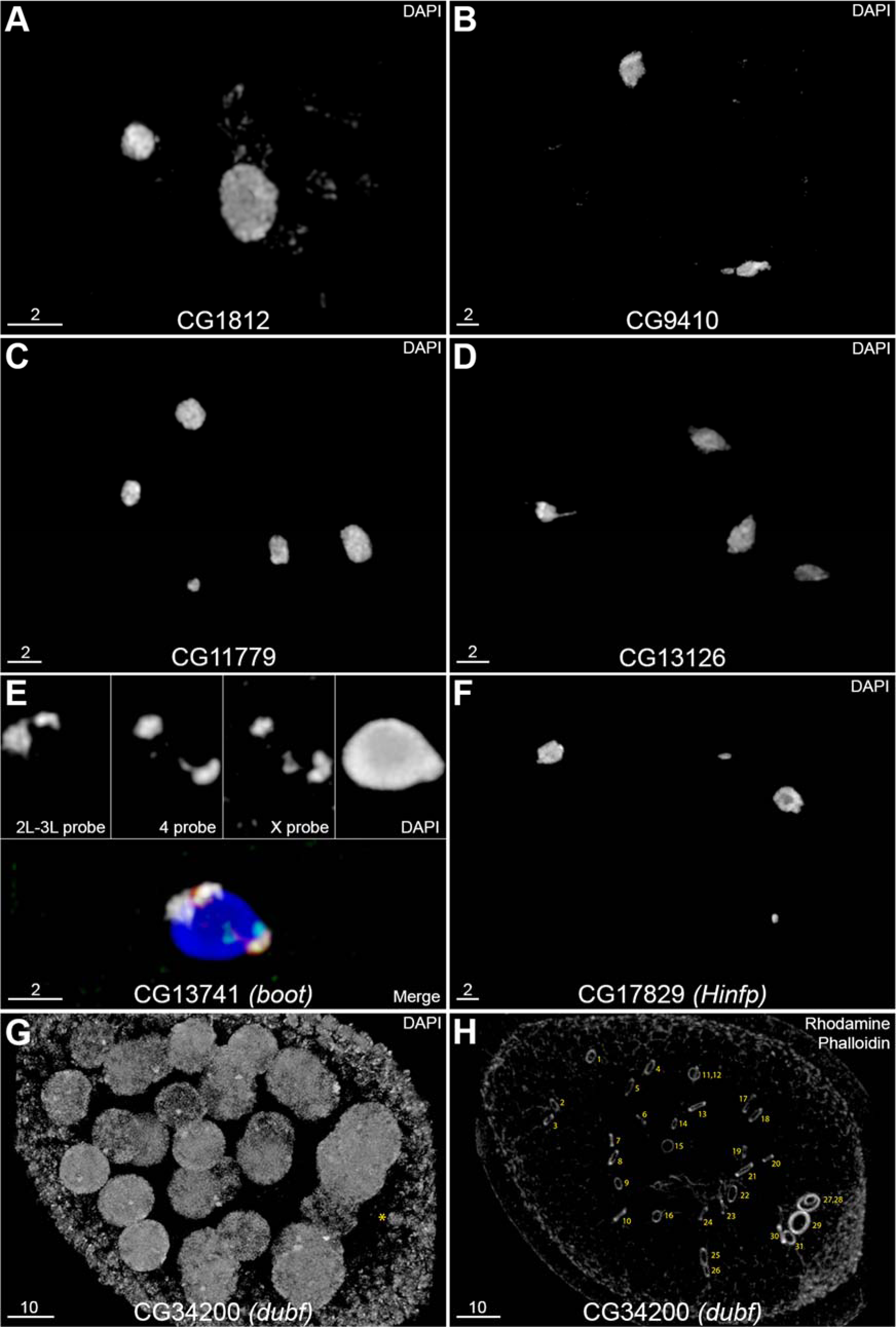
Screen Hits in Previously Uncharacterized Genes. **5A)** RNAi of CG1812 using P{TRiP.GL01163} driven by *matα4::Gal4* resulted in 2 chromosome masses in this oocyte. **5B)** RNAi of CG9410 using P{TRiP.GL01119} driven by *nos::Gal4* resulted in 2 chromosome masses in this oocyte. **5C)** RNAi of CG11779 using the hairpin SH015-H10.N2 driven by *matα4::Gal4* resulted in 5 chromosome masses in this oocyte. This genotype also exhibited nurse cell defects (Figure S1). **5D)** RNAi of CG13126 using P{TRiP.GLC01687} driven by *nos::Gal4* resulted in 4 chromosome masses in this oocyte. **5E)** RNAi of CG13741 *(bootlegger)* using P{TRiP.GL00570} driven by *nos::Gal4* resulted in a failure to coorient all chromosomes. Heterochromatin FISH shows that while the *4* and *X* probes appear to be cooriented on opposite sides of the karyosome, all the probe for *2L-3L* centric heterochromatin is on the same side. Signals from each probe (*2L-3L* = white, *4* = red, *X* = green, DAPI = blue) are shown individually above the false-colored merger. **5F)** RNAi of CG17829 *(Hinfp)* using P{TRiP.GL01162} driven by *nos::Gal4* resulted in 4 chromosome masses in this oocyte. **5G)** RNAi of CG34200 *(dubf)* using P{TRiP.GLC01618} driven by *nos::Gal4* caused this cyst to undergo nurse cell doubling; 31 nurse cells are present in this image, while the yellow asterisk denotes the oocyte nucleus. **5H)** The same CG34200-depleted cyst as 5G, showing rhodamine phalloidin staining of the 31 numbered ring canals. Note that the cell with the oocyte nucleus in the bottom right has 5 rings associated with it, consistent with an additional round of mitosis having occurred. All scale bars are in µm.

#### CG9410

We identified *CG9410* as a *nanos* hit with congression defects, with females undergoing RNAi producing 10/23 oocytes having multiple DNA masses, while no defects in the Mps1-binding filaments were observed (Figure 5B). Ovaries in these females were also very small, with few mature cysts being produced per female. We were able to independently validate this result with a classical allele, as a P element insertion in the 5’ UTR of the CG9410-RA isoform *(P{SUPor-P}CG9410^KG06373^)* was available from the stock center. Hemizygous females carrying that P element over a deletion *(Df(3R)BSC326)* were observed to have a 10% congression failure rate. Females undergoing RNAi were also sterile, whereas *P{SUPor-P}CG9410^KG06373^/Df(3R)BSC326* females were weakly fertile, with >50% of the surviving progeny having been produced from nondisjunctional oocytes (6 normal, 2 *4-*only NDJ, 1 *X-*only NDJ, and 9 *XX⇔44* double NDJ). We also noted that flies carrying the deficiency developed more slowly than wildtype flies, with bottles requiring an additional ∼3 days to start producing progeny. This indicates that *P/Df* is a hypomorphic genotype with meiotic chromosome segregation being seriously compromised by the loss of this gene. According to its entry in Flybase (http://flybase.org/reports/FBgn0033086), *CG9410* has some similarity to coenzyme Q-binding protein *COQ10,* and is predicted to be involved in transporting coenzyme Q6 to the mitochondrial respiratory complexes. Because we were able to validate the phenotype seen by RNAi using an independent reagent, we have given *CG9410* the name *<u>con</u>gression failure in <u>g</u>ermline (config)*.

#### CG11779

We identified *CG11779* as a *mat-alpha* hit, with only 3/20 oocytes having any visible Mps1-binding filaments, and 9/20 oocytes having shredded or completely missing meiotic chromosomes. Oocytes also displayed dorsal appendage and nurse cell deformities (Figure S1). While no mature oocytes were found when RNAi was driven by *nos::Gal4,* females undergoing RNAi driven by *matα4::Gal4* were semi-sterile, with only a few progeny produced per vial. A second-site RNAi construct in VALIUM22 (SH015-H10.N2) was created by the TRiP project, and this construct did recapitulate the phenotype, with 8/13 oocytes having multiple masses as well as serious dorsal appendage and nurse cell defects (Figures 5C and S1). The second allele did not appear to affect the Mps1-binding filaments as strongly, as filaments could be seen in 8/13 oocytes. According to the gene’s entry in FlyBase, *CG11779* is predicted to be orthologous to human *translocase of inner mitochondrial membrane 44 (TIMM44).* Because we were able to validate *CG11779* with an independent RNAi construct, we have named *CG11779* after its human ortholog, *dTIMM44*.

#### CG13126

We identified *CG13126* as a *nanos* hit with congression defects, with females undergoing RNAi producing 8/20 oocytes having multiple DNA masses (Figure 5D), along with a further 5/20 oocytes having shredded or completely missing meiotic chromosomes. No defects in the Mps1-binding filaments were observed. Females undergoing RNAi were also found to be completely sterile. According to the gene’s entry in FlyBase (http://flybase.org/reports/FBgn0032168), the gene is predicted to have methyltransferase activity and is orthologous to the human gene *methyltransferase like 17* (METTL17). As of this writing, creation of a second-site RNAi construct in the VALIUM22 vector by the TRiP project is underway, but as we have not validated this result with a second independent allele, we have not named this gene at this time.

#### CG13741 (Bootlegger)

When we first tested *CG13741* with *nos::Gal4*, we found that chromosomes in all oocytes successfully completed congression to a single mass, while no defect in the Mps1-binding filaments was observed. However, in 5/10 oocytes, we noticed that the brightly-staining heterochromatin in the metaphase-arrested karyosomes formed a single spot, instead of two spots positioned at opposite ends of the karyosome (Fig. 1D). This would suggest that proper chromosome coorientation was not being achieved, as the heterochromatin is normally clustered by the homologous centromeres facing opposite spindle poles. We confirmed this interpretation by chromosome-specific FISH, which showed that homologous chromosomes were maloriented within the karyosome in 11/21 oocytes (Figure 5E). Females undergoing RNAi were sterile, and we have not yet been able to validate this RNAi result using an independent reagent. While we first identified *CG13741* as a hit in 2016, before the publication of this manuscript two groups published linked papers with the first characterization of this gene, showing that it is required for piRNA production, and naming the gene *bootlegger* (ElMaghraby *et al*. 2019; Kneuss *et al*. 2019).

#### CG17829 (Histone Nuclear Factor P)

We identified *CG17829* as a *nanos* hit with congression defects, with 17/30 oocytes having multiple DNA masses, while no defect in the Mps1-binding filaments was observed (Figure 5F). While germline RNAi of this gene with *nos::GAL4* resulted in congression errors, the females were fertile, and a nondisjunction assay showed meiotic nondisjunction occurs at 3.0% for the X and 2.9% for the 4 chromosomes (620 normal progeny, with 17 *4-*only, 9 *X-*only, and 1 *X&4* double nondisjunctional progeny). We have not been able to validate this RNAi result using an independent reagent. While we first identified *CG17829* as a hit in 2018, before the publication of this manuscript another group published the first characterization of this gene, showing that it is required in *Drosophila* for repression of transposable elements in somatic tissues, and identifying it as the homolog of the human gene *histone nuclear factor P (Hinfp)* (Nirala *et al*. 2021).

#### CG34200 (Double Fission)

When *CG34200* was tested, it was found to have no defects in either congression or in the Mps1-binding filaments. However, it was noted that many earlier stage oocytes clearly had more than the usual 15 polytene nurse cells. Because this was an interesting phenotype in a previously uncharacterized gene, we classified it as a *nanos* hit. Counting more cysts showed that more than 15 nurse cells were found in 2/3 of all cysts (28/42), and examination of these cysts with rhodamine phalloidin found that there were 31 nurse cells and 31 ring canals (Figures 5G-H), consistent with one extra round of mitosis with incomplete cytokinesis during cyst development. When RNAi was driven by *matα4::Gal4,* which does not activate until after the nurse cells are established, no nurse cell doubling was observed (0/59 cysts). Despite this, oocytes from RNAi females appeared to complete oogenesis normally, undergoing nurse cell dumping and apoptosis to produce mature eggs that appeared regular sized and were otherwise indistinguishable from wildtype. These females were also fertile, and exhibited normal levels of nondisjunction.

According to this gene’s entry in FlyBase (http://flybase.org/reports/FBgn0085229), this gene is only 52 amino acids long, with no identifiable protein motifs or known human ortholog. This RNAi construct has already been used in a study of ring canal growth rates during oogenesis (L. Llewelyn et al, MS in review, https://doi.org/10.1101/2023.08.18.553876); in that paper *CG34200* was given the name *double fission (dubf)* due to having twice as many nurse cells because of the extra round of division.

### Screen Hits: Genes that Altered Filament Shapes

Several of the screen hits were found to cause noticeable changes to the structure of the Mps1-binding filaments.

#### Mustard

We identified *mustard (mtd)* as a *nanos* hit with both congression defects and changes to the Mps1-GFP binding filaments. Females undergoing RNAi with construct *P{TRiP.GL00598}* driven by *nos::Gal4* had multiple DNA masses in 44 out of 55 oocytes (80%). Those oocytes also displayed a clear change in the Mps1-GFP binding filaments, where instead of individual separated filaments, the filaments appeared to be much longer and formed a more continuous network (Figure 6A). Females undergoing RNAi also exhibited high rates of nondisjunction, with 18.6% X and 9.6% 4 NDJ. When RNAi was driven using *matα4::Gal4*, we saw no congression defects (0 / 30 oocytes), along with normal levels of NDJ (0.2% *X* and 0.1% *4*). As this gene also had a second-site RNAi construct available in the VALIUM22 library (*P{TRiP.GL00665}*) we were able to confirm this screen hit using RNAi. This second allele exhibited congression failure in 43% of oocytes (13/30) and also had the Mps1-GFP binding filaments form a more continuous network (Figure 6A’).

**Figure 6:**
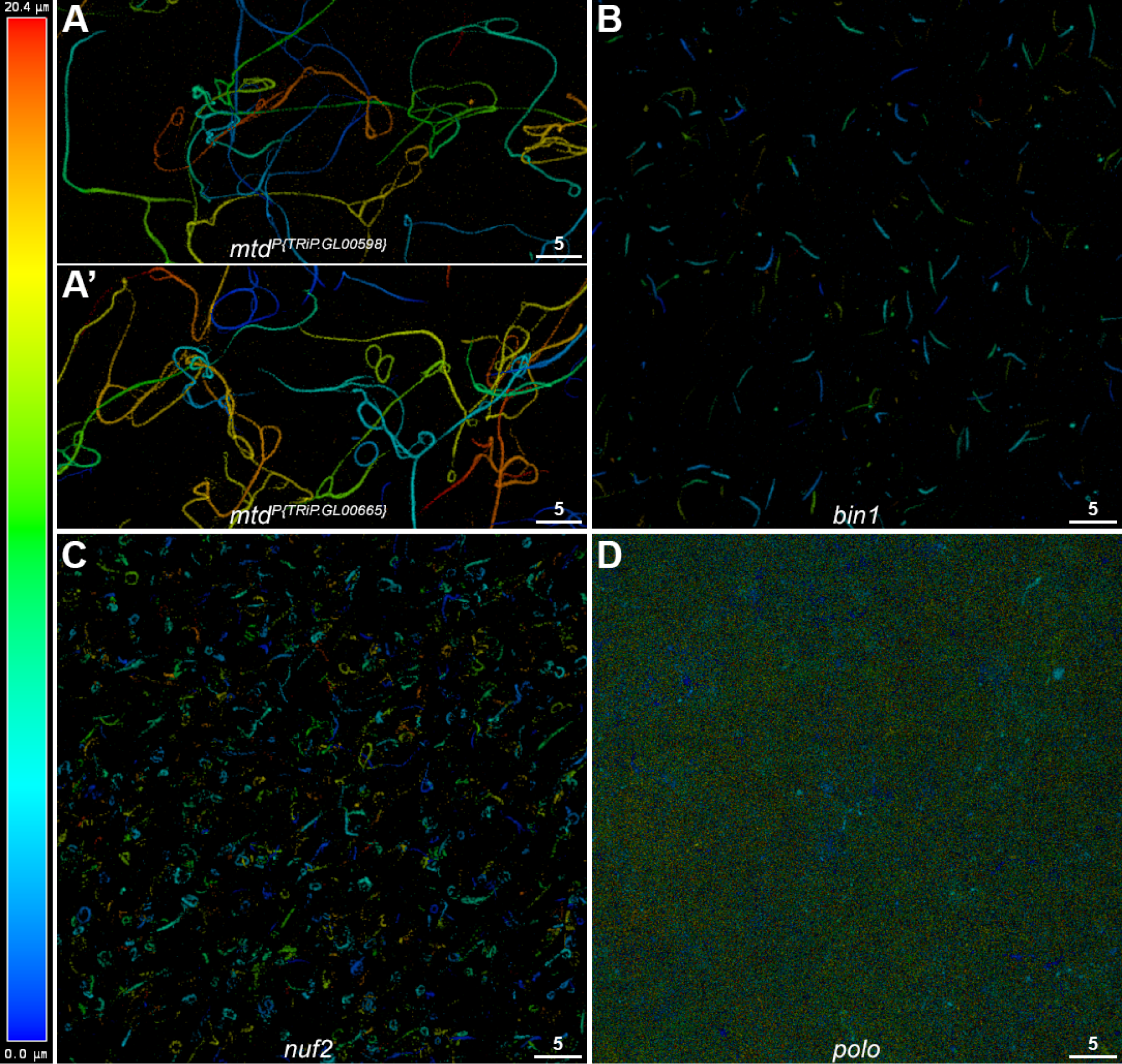
Filament Shape Changes. For all panels, a projection of a 20.4 µm deep Z-stack showing Mps1-GFP fluorescence in a hypoxic oocyte is shown. Imaging depth is indicated according to the false-color scale on the left, with 0 being nearest the objective and positive numbers penetrating deeper into the cell. All scale bars are 5 µm. **6A)** RNAi knockdown of *mtd* using P{TRiP.GL00598} driven by *nos::Gal4*. When compared to Figure 2, filaments are considerably longer and more connected, with fewer isolated individual segments. **6A’)** RNAi knockdown of *mtd* using P{TRiP.GL00665} driven by *nos::Gal4.* This VALIUM22 construct targets a different sequence in *mtd* than P{TRiP.GL00598}, but causes elongated filaments very similar to the other construct, validating this RNAi result. **6B)** RNAi knockdown of *bin1* using P{TRiP.GL00127} driven by *nos::Gal4* results in filaments that are straighter and shorter, mostly consisting of isolated short segments. **6C)** RNAi knockdown of *nuf2* using P{TRiP.GL00436} driven by *nos::Gal4* results in filaments that mostly present as small blobs or tightly-coiled balls instead of long fibers. **6D)** RNAi knockdown of *polo* using P{TRiP.GL00014} driven by *matα4::Gal4* causes most oocytes in this genotype to have no detectable Mps1-binding filaments at all, but approximately 20% of oocytes display very few short filaments along with many small puncta of GFP localization. (Scale bars = 5 µm)

We were further able to confirm that ablation of *mtd* was causing the observed congression defect using a standard mutant allele. A hypomorphic *mtd* allele caused by a P element inserted within the gene (*mtd^KG02600^*) and had been previously reported to cause reduced protein levels (Wang *et al*. 2013) was available from the stock center. We crossed this allele to two different deficiencies that uncovered the *mtd* locus (Df(3R)BSC177 and Df(3R)ED5147), and saw the same 13% rate of congression failures (4/30) in both hemizygous genotypes. While this is a lower rate than the RNAi alleles, it is consistent with this hypomorphic allele having a less compromised *mtd* phenotype than the RNAi treatments. (These genotypes lacked the Mps1-GFP construct, so we did not assess filament shape.)

#### Bin1

We identified *bicoid interacting protein 1 (bin1)* as a *nanos* hit, because while congression appeared to be normal, the Mps1-binding filaments had noticeably different shapes in all oocytes examined. The filaments appeared to be considerably shorter and straighter than in normal flies (Figure 6B). This shape difference was consistently found in repeated dissections. A second-site RNAi construct in VALIUM22 (SH06530.N2) was created by the TRiP project. However, this allele did not recapitulate the phenotype, as filaments did not appear to be shortened relative to wildtype. This may be due to this allele causing less efficient knockdown, but we have not been able to confirm this hypothesis.

#### Nuf2

We identified *nuclear filament-containing protein 2 (nuf2)* as a *nanos* hit, because while congression appeared to be completely normal, when construct SH02141.N2 was driven by *nos::Gal4* many of the Mps1-binding filaments were shaped noticeably differently. Rather than appearing as long thin fibers, the filaments appeared to be twisted up into small balls (Figure 6C). This shape difference was consistently found in repeated dissections. Females with RNAi driven by *nos::Gal4* were weakly fertile, with a rate of *X* nondisjunction over 50% (6 normal progeny, 1 *4-*only and 7 *X*-only nondisjunctional progeny). A second-site RNAi construct in VALIUM22 (SH02715.N2) was created by the TRiP project; however when this construct was driven by *nos::Gal4* it did not cause a similar change in filament shapes (data not shown). However, we believe this is because this construct causes a lower level of knockdown than SH02141.N2, as these females were also more fertile with much lower rates of nondisjunction (56 normal progeny, 1 *4-*only, and 1 *X-*only progeny).

#### Polo

We identified *polo* as a *mat-alpha* hit, as females undergoing RNAi failed to reach a single mass in 5/20 oocytes, and furthermore the Mps1-binding filaments were either completely absent or appeared as a spotty “dust” of foci and small fragments instead of long fibers (Figure 6D). This result extends a previous finding that Polo-GFP protein had been previously demonstrated to be sequestered to the same filaments as Mps1-GFP in response to hypoxia (Gilliland *et al*. 2009b). When RNAi was driven by *nos::Gal4,* females had no stage 14 oocytes, while when driven by *matα4::Gal4,* females were completely sterile. We have not attempted to validate this RNAi result using an independent reagent.

### Screen hits: Other Miscellaneous Results

Two of our hits in previously characterized genes also displayed additional phenotypes that, to the best of our knowledge, have not been previously reported.

#### Lamin

We identified *lam* as a *nanos* hit, as females undergoing RNAi had multiple chromosome masses in 6/10 oocytes, while no defect in the Mps1-binding filaments was observed. However, rather than these multiple DNA masses appearing to be the result of congression failure (which should cause the DNA masses to appear smaller than a normal meiotic karyosome that has all chromosomes properly congressed), these DNA masses appeared to be full-sized meiotic karyosomes (Figure S2), as if the loss of *lam* had caused the cyst to fail to designate a single oocyte nucleus, resulting in multiple nurse cells adopting the oocyte nuclear fate. Lamin is a key component of the nuclear envelope, and is associated with a variety of pathologies, including Hutchison-Gilford Progeria syndrome (Prokocimer *et al*. 2009). We have not attempted to recreate this phenotype with a different reagent.

#### Stem Cell Tumor

We identified *stet* as a *nos* hit, as females undergoing RNAi had multiple chromosome masses in 5/20 oocytes, while no defect in the Mps1-binding filaments was observed. Females undergoing RNAi were semi-sterile. Interestingly, in addition to a moderate level of nurse cell doubling seen in prophase cysts, all of the mature eggs produced by these females failed to develop any dorsal appendages (Figure S2). The *stet* gene is a homolog of *rhomboid* that activates ligands of the Egfr signaling pathway, and is believed to regulate germ cell encapsulation by somatic cells (Schulz *et al*. 2002). We have not attempted to recreate this phenotype with a different reagent.

## Discussion

One of the motivations for this project was the prediction that many genes of interest to the study of meiosis could never be recovered in traditional genetic screens, as the loss of that gene would result in either sterility or lethality in the mother and make it impossible to look for segregation errors in her progeny. However, by restricting RNAi knockdown to the germline, we were able to test genes that would be required for viability if ablated in all tissues, and by examining unfertilized eggs, we were able to assay sterile genotypes. That this motivation was well founded is demonstrated by the fact that over two-thirds of our hits (35/52 tested constructs) were sterile or semi-sterile when undergoing RNAi, while less than one third (17/52) were fertile. As three of the novel genes that we are the first to associate phenotypes with were sterile (*CG1812, CG11779, and CG13126*), we believe that our results clearly demonstrate that previous genetic screens that measured nondisjunction missed many of the genes required for female meiosis in *Drosophila*.

While our use of RNAi did provide several advantages for the screen, we did run into a number of the limitations inherent to the technique. RNAi requires validation of positive results, and we had hoped to use stocks from the larger VALIUM20 collection for validation. However, none of the constructs that we tested from that library (0/8) showed any phenotypes, presumably because that vector has a lower level of germline transcription when compared to VALIUM22. As many more second-site constructs were available from the VALIUM20 collection, this limited the reagents available to confirm screen results. However, even within VALIUM22, only three of the second-site constructs we tested validated the original phenotypes (*cdk2, CG11779, mtd*), while four others did not (*bin1, CG1812, krimp, nuf2*). This under-50% validation rate suggests that a substantial number of constructs likely yielded false negative results, and would have been classified as hits if the knockdown could have been stronger.

It is important to remember that a second-site construct’s failure to confirm is a negative inference, and does not invalidate the original positive result. We have not yet pursued why these second-site alleles failed, but it seems reasonable that the level of knockdown should correspond to variation in the phenotypes we saw. For example, the original *nuf2* allele (P{TRiP.GL00436}) caused a significant filament shape defect (Figure 6C) and was only weakly fertile, with high levels of nondisjunction. A new VALIUM22 hairpin (SH02715.N2) was created by the TRiP project, but in addition to not displaying the same filament shape defect, it also had better fertility and a much lower nondisjunction rate. This would suggest the new RNAi construct is simply not causing sufficient knockdown of Nuf2 to cause the phenotypes seen in the original allele. This explanation can be tested directly by qRT-PCR or southern blot, by driving expression of SH02715.N2 more strongly by using the stronger Maternal Triple Driver stock or by including additional copies of the *nos::Gal4* driver construct, or by including a deficiency that removes one of the endogenous copies of the *nuf2* locus, which should reduce the amount of transcript that RNAi needs to eliminate.

Because the screen identified several different sets of genes (hits, *nos* duds, and untestable genes) we performed a Gene Ontology analysis, to see what biological processes were most represented within each group. One unexpected result of this analysis was the abundance of genes involved in RNA metabolism present in both the *nos* duds and the screen hits. Among the hits, nearly all of these RNA process genes (22/24) only caused chromosome congression errors without affecting the filaments, while the two exceptions *(cuff, wisp)* were previously known meiotic genes that cause major defects in oogenesis (Cui *et al*. 2008; Mohn *et al*. 2014). Curiously, *cuff* and *wisp* affected the Mps1-binding filaments in opposite directions, with RNAi of *cuff* making the filaments appear to be brighter than normal, while RNAi of *wisp* caused the filaments to be absent. As both genes regulate maternal mRNA transcripts (Chen *et al*. 2007; Dufourt *et al*. 2017), this raises the possibility that one function of the Mps1-binding filaments may be to regulate mRNAs during hypoxia. In fact, out of the ∼30 genes known to be involved in piRNA biogenesis, five of them (*cuff, wisp, boot, csul, tej)* were among our hits.

While only 8/24 of the RNA process genes had been previously recognized as required for meiosis, this result is concordant with several recent findings that implicate RNA metabolic functions as being important for the establishment of homologous chromosome pairing, which might explain the congression phenotypes we observed. In studies done in the fission yeast *S. pombe,* the process of homologous pairing was found to depend on RNA-protein complexes (Ding *et al*. 2019), as well as long non-coding RNAs being required for the transition from mitosis to meiosis (Andric *et al*. 2021). The processes of both transcription and splicing have also been implicated in establishing the regular interchromosomal chromatin architecture in fission yeast (Bertero 2021). In addition to findings in yeast, a study in *D. melanogaster* found that pairing during early embryogenesis occurred at the same time point that zygotic transcription is activated (Erceg *et al*. 2019). As proper homologous pairing is required for meiotic recombination in *Drosophila* (Takeo *et al*. 2011), disruption of RNA metabolism could therefore disrupt a pairing process, which would in turn inhibit recombination, and that is well known to cause meiotic nondisjunction (Hawley *et al*. 1993). Therefore, it appears that our results are consistent with this developing story, and that knocking down different components of the RNA metabolism in developing oocytes may lead to congression failure. We note that among the RNA-associated hits that were fertile, most (5/8) also had elevated NDJ rates. This model makes a simple prediction: when RNAi of RNA-associated genes is driven with *nos::Gal4,* we should observe reduced recombination rates in the progeny, but not when RNAi is driven by *mata::Gal4*, as that driver does not start expressing until after recombination is completed. (However, we also note that recombination cannot be the sole cause of this congression failure, as 9/25 of the RNA-associated hits were driven by *mata::Gal4*.) Our lists of hits and *nos* duds should provide useful candidate genes for further study of the roles that RNA-related genes play during female meiosis.

One major goal of this project was to identify one or more of the major structural protein components that form the scaffold of the Mps1-binding filaments. While we did not achieve this goal, we did identify several genes that clearly affected the filaments. Several genes known to be required for meiosis, such as *cdk1, twe,* and *wisp,* caused the filaments to be completely absent in many oocytes. As the filaments appear to form at germinal vesicle breakdown (GVBD) (Gilliland *et al*. 2009b) and disappear shortly after egg activation (Pandey *et al*. 2007), this would be consistent with disruption of these genes causing the filaments to not be present at their usual time during prometaphase. In addition, several genes were found to alter the shapes of the filaments. RNAi of *bin1 (Bicoid INteracting protein 1)* caused the filaments to appear shorter and straighter than normal (Figure 6B) while RNAi of *nuf2 (NUclear Filament-containing protein 2)* caused the filaments to appear like twisty balls instead of long strings (Figure 6C). Both genes appear to be very relevant, given what little is known about the filaments. The filaments start to assemble at GVBD, and appear to do so in a wave that progresses from the posterior to the anterior end of the oocyte (Gilliland *et al*. 2009b). The posterior-anterior axis of the oocyte is established by a gradient of Bicoid protein (Driever and Nusslein-Volhard 1988), which suggests that the change in filament shape caused by *bin1* RNAi may reflect the way that the filaments are assembled. Likewise, the observation of filament changes by *nuf2* RNAi is interesting, as the *C. elegans* homolog of *nuf2, HIM-10,* was one of the proteins that was found localized to very similar appearing ooplasmic filaments found in nematode oocytes (Monen *et al*. 2005). While it is not known whether that filament localization in *C. elegans* is also a dynamic response to hypoxia, it suggests that the ooplasmic filaments may be conserved structures that predate the divergence of arthropods and nematodes.

We also note that depletion of *mtd* causes the filaments to appear as a more connected network instead of independent segments (Figures 6A-B). This suggests a hypothesis that the underlying scaffold proteins are normally in that more continuous network, and that one function of Mtd is to establish domains in the network where Mps1-GFP binding is excluded, while other domains still can bind it. However, without Mtd the exclusionary domains are never established, and the entire filament network can bind to Mps1-GFP, resulting in the more continuous filament network that is observed in *mtd-*RNAi females. This model predicts that Mtd protein should bind to the filaments in an alternating fashion with Mps1. We note that the UniProt database (entry A0A0B4K620) lists Mtd as having a LysM domain, which can bind structural molecules such as peptidoglycans (Mesnage *et al*. 2014), suggesting that class of structural molecules as possible future candidates for the structural components of the ooplasmic Mps1-binding filaments.

Going into this project, our naïve assumption was that the Mps1-binding filaments were dispensable, and that we would be able to knock them out completely and still have mature stage 14 eggs to score. However, all of the genes that we found that changed the filament shapes *(bin1, mtd,* and *nuf2)*, along with the genes known to localize to the filaments *(mps1* and *polo),* have lethal alleles. This suggests that the Mps1-binding scaffolds may actually be essential structures, and our failure to identify their structural components is because it is simply not possible to complete oogenesis without them. While this outcome would certainly be ironic, given our motivations for screening by RNAi in the first place, it does suggest that the structural components of the scaffold that we have been seeking may yet be found within the lists of duds and untestable genes.

In conclusion, this project was an ambitious attempt to do a large-scale genetic screen at a primarily undergraduate institution. Despite the limitations inherent to RNA interference, we identified over 50 hits, some of which we have already successfully validated using additional reagents. Going forward, the list of hits will provide fruitful projects for students to continue working on for years to come. These future projects include more focused validation attempts (e.g. by enhancing RNAi knockdown via including additional copies of the Gal4 driver or deficiencies that reduce the level of endogenous RNA, by characterization of other mutant alleles, and by creating RNAi-resistant alleles via CRISPr-induced silent mutations in the regions bound by shRNA hairpins) as well as investigating questions raised by the screen results, such as if knocking down RNA-process genes leads to a reduction in recombination rates, or whether *mtd* causes regions of the ooplasmic filaments to not bind Mps1-GFP.

## Acknowledgements

This work was supported by NIH/NIGMS grant R15GM099054 to WDG. We thank the Bloomington Drosophila Stock Center (NIH P40OD018537) for the many stocks used in this study, as well as Dr. Jonathan Zirin and the TRiP project at Harvard Medical School (NIH/NIGMS R01-GM084947) for both creating the transgenic RNAi fly collections used in this study, and for producing several new VALIUM22 constructs to attempt to confirm some of our hits. We also thank Dr. Justin Blumenstiel, Dr. Margaret Silliker, and Dr. Jeff Sekelsky for helpful comments on the manuscript.

---

**File S1: RNAi Stock Lists**

An Excel table with four sheets, each presenting a supplemental table. **Table S1:** all 1265 negative result lines, which produced scorable oocytes when crossed to NSS, but were not found to exhibit either screened-for phenotype, **Table S2:** all 32 untestable lines, which did not produce scorable oocytes when crossed to either NSS or MASS, **Table S3:** all 108 *nos* dud lines, which did not produce scorable oocytes when crossed to NSS, but were found to have negative results when crossed to MASS, and **Table S4:** construct information for all 54 screen hits, including the second-site VALIUM22 constructs that were tested to validate hits. Tables S1-S3 are sorted by their BDSC stock numbers, while Table S4 is sorted by gene name. All columns except “Status” and “Driver” were copied from the original lists of constructs downloaded from TRiP, and may have been updated in the live TRiP database.

**File S2: ShinyGO outputs**

This Excel table with four sheets, presenting (1) the lists of FBgn identifiers used for the Gene Ontology analyses, with (2-4) being the Groups outputs from the ShinyGO runs that produced Figures 4A-C, identifying which genes are associated with each GO term.

**Figure S1:**
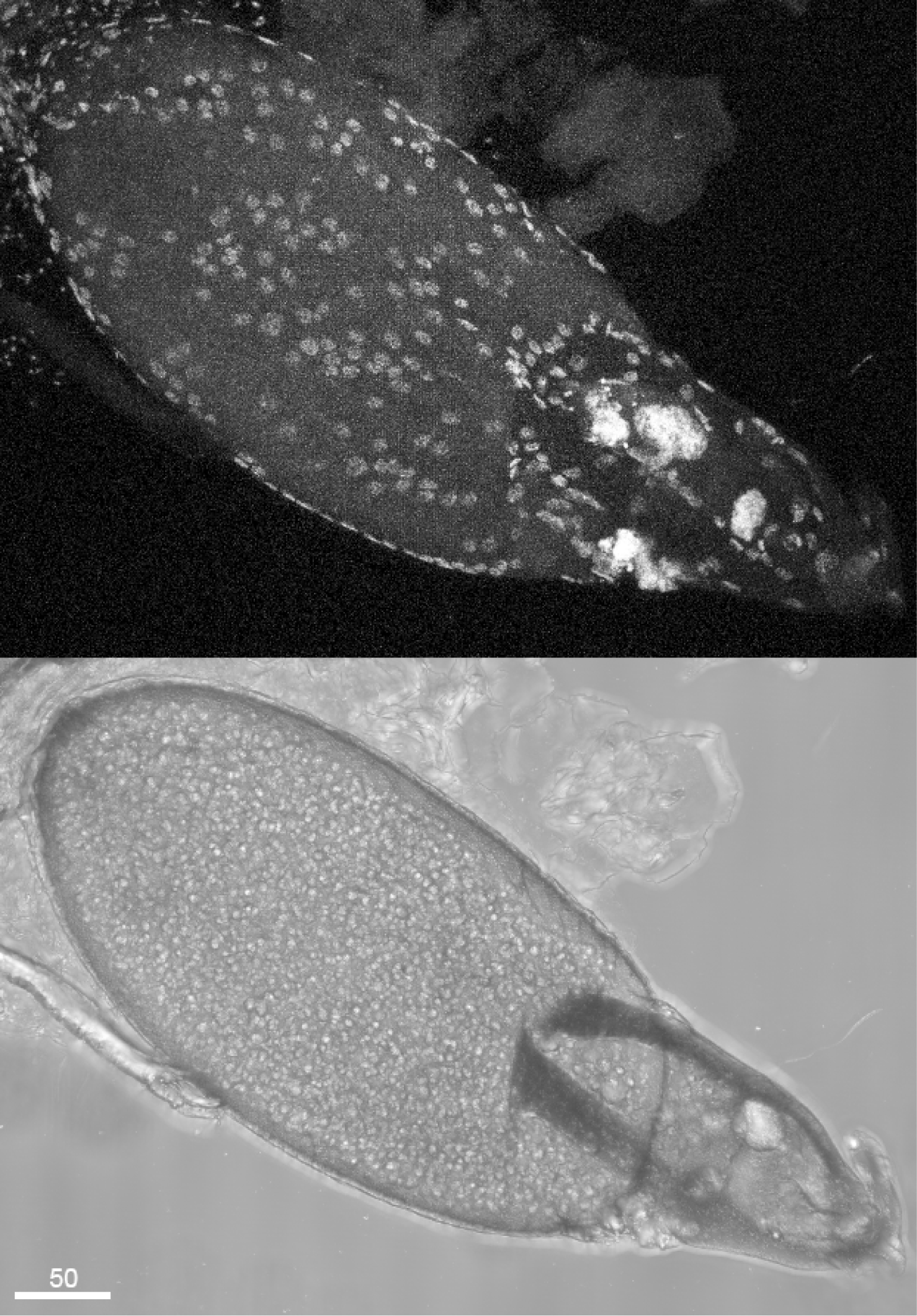
Nurse cell defects in *CG11779*. A low magnification image of the same *CG11779* oocyte used in Figure 5C, with DAPI staining (top) and brightfield (bottom). The 5 DNA masses in Figure 5C are not discernable at this magnification, as they are obscured by polytene follicle cells near the base of the dorsal appendages. This shows the typical nurse cell defect seen in this genotype, where despite having well grown dorsal appendages, the nurse cells did not complete apoptosis and contain material that was not transported into the oocyte. (Scale Bar = 50 µm)

**Figure S2:**
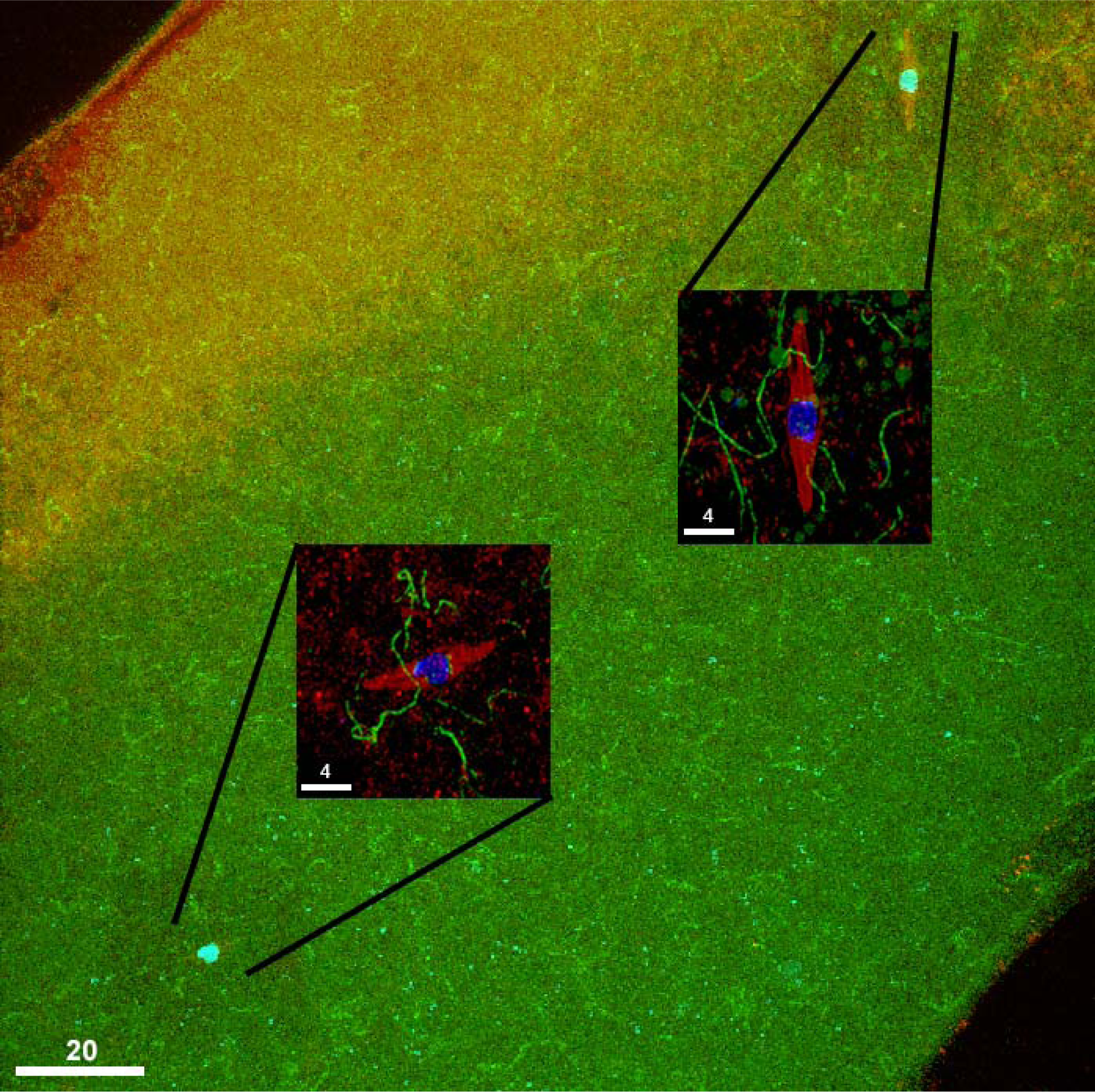
Knockdown of *lamin* Produces Oocytes With Multiple Pronuclei. A low magnification immunofluorescent image of an oocyte undergoing RNAi for *lam,* stained with DAPI (blue), anti-tubulin antibody (red), and Mps1-GFP (green), showing that there are multiple oocyte nuclei present. The sides of the oocyte are visible in the upper left and lower right; the site where the dorsal appendages were is just out of frame in the upper right. The two insets are higher magnification images of the two karyosomes, each with their own tubulin spindle. Each individual DNA mass is the same length and width of a normal meiotic karyosome, and the Mps1-GFP is localized to more than 8 total kinetochore foci, indicating that this represents the formation of multiple oocyte nuclei instead of a congression failure. (Scale Bar = 20 µm)

**Figure S3:**
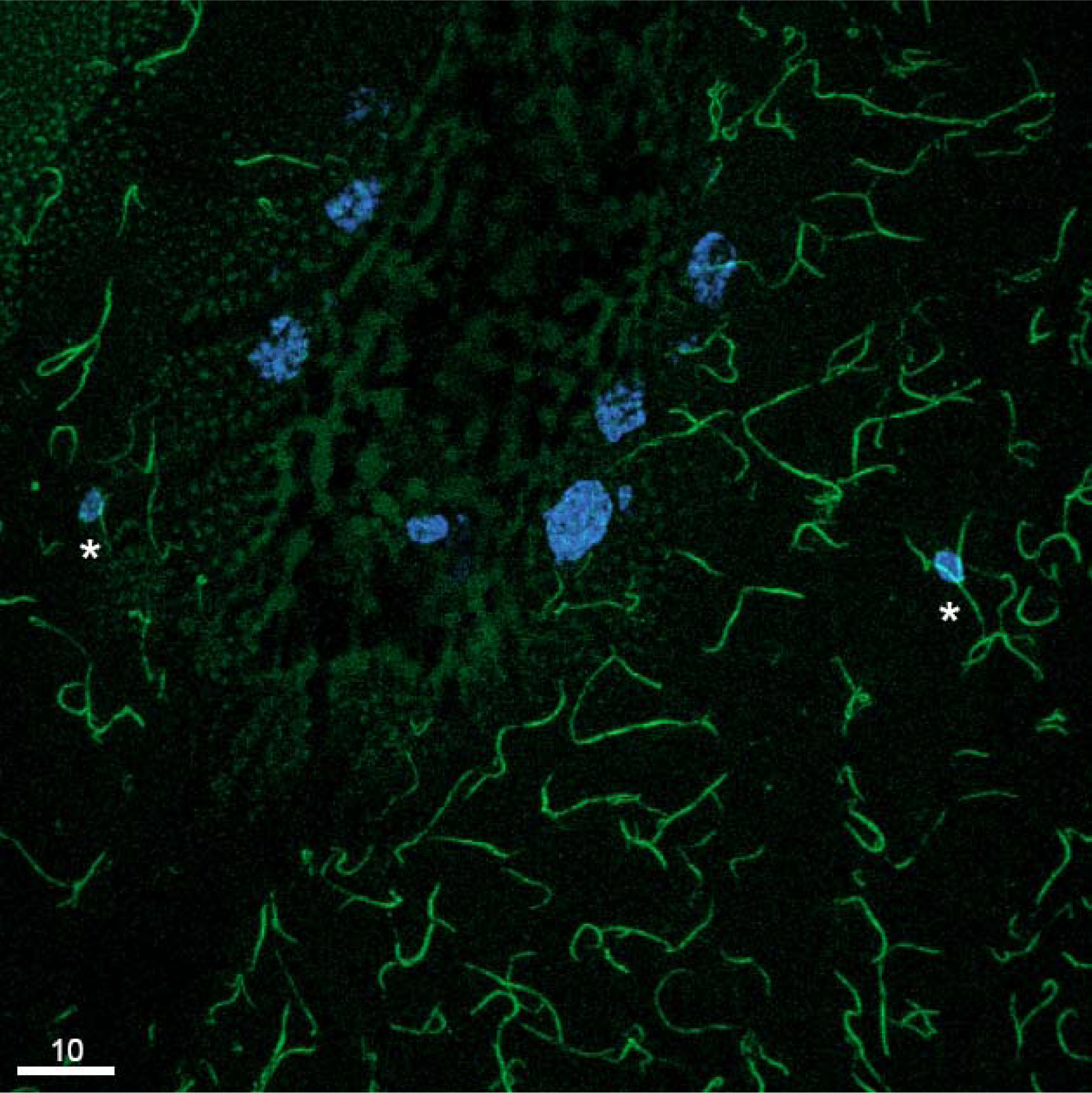
Knockdown of *stem cell tumor* Disrupts Dorsal Appendage Development. An oocyte undergoing RNAi of *stet,* showing DNA (DAPI, blue) and Mps1-GFP (green). This oocyte had two DNA masses (asterisks) on opposite sides of the anterior region of the oocyte where the dorsal appendages should have formed, which retains a few polytene nurse cells that did not complete apoptosis. The GFP signal also detects autofluorescence from the oocyte cortex (the stippled dots in the upper left corner and the fainter blobs around the nurse cells). No mature oocytes were found to develop dorsal appendages in this genotype.

## Bibliography

Andric V., A. Nevers, D. Hazra, S. Auxilien, A. Menant, et al., 2021 A scaffold lncRNA shapes the mitosis to meiosis switch. Nat Commun 12: 770. 10.1038/s41467-021-21032-7

Baker B. S., and A. T. Carpenter, 1972 Genetic analysis of sex chromosomal meiotic mutants in Drosophilia melanogaster. Genetics 71: 255–86. 10.1093/genetics/71.2.255

Bertero A., 2021 RNA Biogenesis Instructs Functional Inter-Chromosomal Genome Architecture. Front Genet 12: 645863. 10.3389/fgene.2021.645863

Chen Y., A. Pane, and T. Schüpbach, 2007 cutoff and aubergine Mutations Result in Retrotransposon Upregulation and Checkpoint Activation in Drosophila. Current Biology 17: 637–642. 10.1016/j.cub.2007.02.027

Cook K. R., A. L. Parks, L. M. Jacobus, T. C. Kaufman, and K. A. Matthews, 2010 New research resources at the Bloomington Drosophila Stock Center. Fly (Austin) 4: 88–91. 10.4161/fly.4.1.11230

Cooper K. W., 1948 A New Theory of Secondary Non-Disjunction in Female Drosophila Melanogaster. Proc Natl Acad Sci U S A 34: 179–187. 10.1073/pnas.34.5.179

Cui J., K. L. Sackton, V. L. Horner, K. E. Kumar, and M. F. Wolfner, 2008 Wispy, the Drosophila homolog of GLD-2, is required during oogenesis and egg activation. Genetics 178: 2017–29. 10.1534/genetics.107.084558

Ding D. Q., K. Okamasa, Y. Katou, E. Oya, J. I. Nakayama, et al., 2019 Chromosome-associated RNA-protein complexes promote pairing of homologous chromosomes during meiosis in Schizosaccharomyces pombe. Nat Commun 10: 5598. 10.1038/s41467-019-13609-0

Driever W., and C. Nusslein-Volhard, 1988 The bicoid protein determines position in the Drosophila embryo in a concentration-dependent manner. Cell 54: 95–104. 10.1016/0092-8674(88)90183-3

Dufourt J., G. Bontonou, A. Chartier, C. Jahan, A.-C. Meunier, et al., 2017 piRNAs and Aubergine cooperate with Wispy poly(A) polymerase to stabilize mRNAs in the germ plasm. Nat Commun 8: 1305. 10.1038/s41467-017-01431-5

ElMaghraby M. F., P. R. Andersen, F. Pühringer, U. Hohmann, K. Meixner, et al., 2019 A Heterochromatin-Specific RNA Export Pathway Facilitates piRNA Production. Cell 178: 964–979.e20. 10.1016/j.cell.2019.07.007

Erceg J., J. AlHaj Abed, A. Goloborodko, B. R. Lajoie, G. Fudenberg, et al., 2019 The genome-wide multi-layered architecture of chromosome pairing in early Drosophila embryos. Nat Commun 10: 4486. 10.1038/s41467-019-12211-8

Fischer M. G., S. Heeger, U. Hacker, and C. F. Lehner, 2004 The mitotic arrest in response to hypoxia and of polar bodies during early embryogenesis requires Drosophila Mps1. Curr Biol 14: 2019–24. 10.1016/j.cub.2004.11.008

Ge S. X., D. Jung, and R. Yao, 2020 ShinyGO: a graphical gene-set enrichment tool for animals and plants. Bioinformatics 36: 2628–2629. 10.1093/bioinformatics/btz931

Gillies S. C., F. M. Lane, W. Paik, K. Pyrtel, N. T. Wallace, et al., 2013 Nondisjunctional segregations in Drosophila female meiosis I are preceded by homolog malorientation at metaphase arrest. Genetics 193: 443–51. 10.1534/genetics.112.146241

Gilliland W. D., S. E. Hughes, J. L. Cotitta, S. Takeo, Y. Xiang, et al., 2007 The multiple roles of mps1 in Drosophila female meiosis. PLoS Genet 3: e113. 10.1371/journal.pgen.0030113

Gilliland W. D., S. F. Hughes, D. R. Vietti, and R. S. Hawley, 2009a Congression of achiasmate chromosomes to the metaphase plate in Drosophila melanogaster oocytes. Dev Biol 325: 122–8. 10.1016/j.ydbio.2008.10.003

Gilliland W. D., D. L. Vietti, N. M. Schweppe, F. Guo, T. J. Johnson, et al., 2009b Hypoxia transiently sequesters mps1 and polo to collagenase-sensitive filaments in Drosophila prometaphase oocytes. PLoS One 4: e7544. 10.1371/journal.pone.0007544

Gilliland W. D., E. M. Colwell, F. M. Lane, and A. A. Snouffer, 2014 Behavior of aberrant chromosome configurations in Drosophila melanogaster female meiosis I. G3 (Bethesda) 5: 175–82. 10.1534/g3.114.014316

Hawley R. S., K. S. McKim, and T. Arbel, 1993 MEIOTIC SEGREGATION IN *DROSOPHILA MELANOGASTER* FEMALES: MOLECULES, MECHANISMS, AND MYTHS. Annu. Rev. Genet. 27: 281–319. 10.1146/annurev.ge.27.120193.001433

Heigwer F., F. Port, and M. Boutros, 2018 RNA Interference (RNAi) Screening in Drosophila. Genetics 208: 853–874. 10.1534/genetics.117.300077

Hughes S. E., W. D. Gilliland, J. L. Cotitta, S. Takeo, K. A. Collins, et al., 2009 Heterochromatic threads connect oscillating chromosomes during prometaphase I in Drosophila oocytes. PLoS Genet 5: e1000348. 10.1371/journal.pgen.1000348

King R. C., 1970 Ovarian development in Drosophila melanogaster. Academic Press, New York,.

Kneuss E., M. Munafò, E. L. Eastwood, U.-S. Deumer, J. B. Preall, et al., 2019 Specialization of the Drosophila nuclear export family protein Nxf3 for piRNA precursor export. Genes Dev. 33: 1208–1220. 10.1101/gad.328690.119

Marygold S. J., P. C. Leyland, R. L. Seal, J. L. Goodman, J. Thurmond, et al., 2013 FlyBase: improvements to the bibliography. Nucleic Acids Res 41: D751–7. 10.1093/nar/gks1024

Mazzalupo S., and L. Cooley, 2006 Illuminating the role of caspases during Drosophila oogenesis. Cell Death Differ 13: 1950–9. 10.1038/sj.cdd.4401892

Mesnage S., M. Dellarole, N. J. Baxter, J. B. Rouget, J. D. Dimitrov, et al., 2014 Molecular basis for bacterial peptidoglycan recognition by LysM domains. Nat Commun 5: 4269. 10.1038/ncomms5269

Mohn F., G. Sienski, D. Handler, and J. Brennecke, 2014 The rhino-deadlock-cutoff complex licenses noncanonical transcription of dual-strand piRNA clusters in Drosophila. Cell 157: 1364–1379. 10.1016/j.cell.2014.04.031

Monen J., P. S. Maddox, F. Hyndman, K. Oegema, and A. Desai, 2005 Differential role of CENP-A in the segregation of holocentric C. elegans chromosomes during meiosis and mitosis. Nat Cell Biol 7: 1248–55. 10.1038/ncb1331

Ni J. Q., R. Zhou, B. Czech, L. P. Liu, L. Holderbaum, et al., 2011 A genome-scale shRNA resource for transgenic RNAi in Drosophila. Nat Methods 8: 405–7. 10.1038/nmeth.1592

Nirala N. K., Q. Li, P. N. Ghule, H. J. Chen, R. Li, et al., 2021 Hinfp is a guardian of the somatic genome by repressing transposable elements. Proc Natl Acad Sci U S A 118. 10.1073/pnas.2100839118

Page S. L., R. J. Nielsen, K. Teeter, C. M. Lake, S. Ong, et al., 2007 A germline clone screen for meiotic mutants in Drosophila melanogaster. Fly (Austin) 1: 172–81. 10.4161/fly.4720

Pandey R., S. Heeger, and C. F. Lehner, 2007 Rapid effects of acute anoxia on spindle kinetochore interactions activate the mitotic spindle checkpoint. J Cell Sci 120: 2807–18. 10.1242/jcs.007690

Prokocimer M., M. Davidovich, M. Nissim-Rafinia, N. Wiesel-Motiuk, D. Z. Bar, et al., 2009 Nuclear lamins: key regulators of nuclear structure and activities. J Cell Mol Med 13: 1059–1085. 10.1111/j.1582-4934.2008.00676.x

Rørth P., 1998 Gal4 in the Drosophila female germline. Mechanisms of Development 78: 113–118. 10.1016/s0925-4773(98)00157-9

Sandler L., D. L. Lindsley, B. Nicoletti, and G. Trippa, 1968 Mutants affecting meiosis in natural populations of Drosophila melanogaster. Genetics 60: 525–58. 10.1093/genetics/60.3.525

Schulz C., C. G. Wood, D. L. Jones, S. I. Tazuke, and M. T. Fuller, 2002 Signaling from germ cells mediated by the *rhomboid* homolog *stet* organizes encapsulation by somatic support cells. Development 129: 4523–4534. 10.1242/dev.129.19.4523

Sekelsky J. J., K. S. McKim, L. Messina, R. L. French, W. D. Hurley, et al., 1999 Identification of novel Drosophila meiotic genes recovered in a P-element screen. Genetics 152: 529–42. 10.1093/genetics/152.2.529

Sullivan W., M. Ashburner, and R. S. Hawley, 2000 Drosophila protocols. Cold Spring Harbor Laboratory Press, Cold Spring Harbor, N.Y.

Takeo S., C. M. Lake, E. Morais-de-Sa, C. E. Sunkel, and R. S. Hawley, 2011 Synaptonemal complex-dependent centromeric clustering and the initiation of synapsis in Drosophila oocytes. Curr Biol 21: 1845–51. 10.1016/j.cub.2011.09.044

Theurkauf W. E., and R. S. Hawley, 1992 Meiotic spindle assembly in Drosophila females: behavior of nonexchange chromosomes and the effects of mutations in the nod kinesin-like protein. J Cell Biol 116: 1167–80. 10.1083/jcb.116.5.1167

Wang Z., S. Hang, A. E. Purdy, and P. I. Watnick, 2013 Mutations in the IMD pathway and mustard counter Vibrio cholerae suppression of intestinal stem cell division in Drosophila. mBio 4: e00337–13. 10.1128/mBio.00337-13

Yan D., R. A. Neumuller, M. Buckner, K. Ayers, H. Li, et al., 2014 A regulatory network of Drosophila germline stem cell self-renewal. Dev Cell 28: 459–73. 10.1016/j.devcel.2014.01.020

Zeng Y., H. Li, N. M. Schweppe, R. S. Hawley, and W. D. Gilliland, 2010 Statistical analysis of nondisjunction assays in Drosophila. Genetics 186: 505–13. 10.1534/genetics.110.118778

Zhang P., and R. S. Hawley, 1990 The genetic analysis of distributive segregation in Drosophila melanogaster. II. Further genetic analysis of the nod locus. Genetics 125: 115–27. 10.1093/genetics/125.1.115

Zwick M. E., J. L. Salstrom, and C. H. Langley, 1999 Genetic variation in rates of nondisjunction: association of two naturally occurring polymorphisms in the chromokinesin nod with increased rates of nondisjunction in Drosophila melanogaster. Genetics 152: 1605–14. 10.1093/genetics/152.4.1605

